# Longitudinal spatial profiling of neutrophils during adoptive T cell therapy in murine melanoma reveals distinct lymph node infiltration patterns across anatomical sites

**DOI:** 10.64898/2026.02.24.707484

**Authors:** Gemma van der Voort, Maike Effern, Michelle C. R. Yong, Lukas Kiwitz, Roberta Turiello, Sonia Leonardelli, Susanna Ng, Dillon Corvino, Tobias Bald, Nicole Glodde, Kevin Thurley, Jan Hasenauer, Michael Hölzel

## Abstract

Reactive neutrophil infiltration can restrain CD8^+^ T cell expansion in lymph nodes during adoptive T cell therapy (ACT), yet its spatiotemporal regulation remains incompletely understood. Levaraging flow cytometry and multiplex immunofluorescence data, we performed a time-resolved quantitative assessment of immune cell dynamics in tumor-draining lymph node (tdLN) and non-tumor-draining lymph node (non-tdLN) in a melanoma mouse model receiving ACT. Transferred tumor-reactive CD8^+^ T cells accumulated and expanded early after treatment initiation, showing the highest frequency of a favorable central memory CD8^+^ T cell phenotype in the tdLN. Enhancing innate immune signaling in melanomas increased neutrophil influx into lymph nodes, particularly the non-tdLN; however, within the tdLN, neutrophils were enriched in the T cell zone, which also contained the largest absolute reservoir of transferred CD8^+^ T cells. Together, these findings indicate that tdLN and non-tdLN differ in early neutrophil dynamics and compartmentalization during ACT, influenced by the strength of innate immune signaling in the tumor.

## Introduction

Adoptive T cell therapy (ACT) is a promising strategy for cancer treatment not only in hematological malignancies, but progressively also in solid tumors [1] such as melanoma [2, 3]. Among the different ACT strategies, autologous, *ex vivo* expanded tumor-infiltrating lymphocytes (TILs) have shown promise for melanoma patients. In a recent phase III clinical trial among patients with advanced melanoma refractory to anti-PD-1 therapy, median progression-free survival increased from 3.1 months in the group receiving ipilimumab immune-checkpoint inhibition to 7.2 months in the TIL group [4].

Among the TILs, the CD8^+^ T cells are the primary effector cells mediating therapeutic efficacy, and their abundance is predictive of improved treatment outcomes in a variety of cancer types and treatment regimens [5]. For a sustained CD8^+^ T cell response, maintaining a long-term proliferative capacity is essential [6, 7]. The tumor-draining lymph node (tdLN) serves as a reservoir of stem-like CD8^+^ T cells with strong proliferative capacity [8]. It is an essential site for anti-tumor immune coordination and surveillance [9]. Stem-like CD8^+^ T cells are initially activated in the lymph node and then migrate to the tumor to differentiate into their terminal effector state [10].

The tdLN undergoes significant changes during tumor progression. In its role as the primary site of antigen presentation and T cell activation, this lymph node is exposed to an influx of antigen-presenting cells and undergoes remodeling of its B and T cell compartments [9, 11, 12]. Due to its constant exchange with the tumor, the tdLN is additionally exposed to tumor-mediated immune microenvironment remodeling [13, 14]. Together, these effects result in a size increase of the tdLN that is observed in both human patients and mouse models of cancer [15–17].

In the tdLN, neutrophils are emerging as an important player in tumor-immune dynamics [18, 19]. Traditionally, neutrophils have been thought of as short-lived effector cells that patrol the blood and rapidly enter damaged or infected tissues [20]. Over the last two decades, their more complex roles in cancer and in shaping the immune environment of lymph nodes have become increasingly apparent. Neutrophils play a dual role in cancer, anti-tumoral in early and pro-tumoral in late stages [18, 19, 21]. In the tumor micro-environment (TME), tumor-associated neutrophils or polymorphonuclear myeloid-derived suppressor cells are associated with reduced T cell function and worsening treatment outcomes in many solid cancers [22–24]. Neutrophils are known to interact with CD8^+^ T cells in the TME of various cancers and are recognized as modulators of T cell function during ACT [25, 26]. These interactions can lead to reduced fitness and an impaired anti-tumor response [27–29]. While present in small numbers during homeostasis [30], neutrophils infiltrate both draining and non-draining lymph nodes during cancer progression [31]. In the tdLN, neutrophils can acquire an immunosuppressive phenotype similar to their counterparts in the TME, as shown in samples from patients with head and neck cancer (HNC) [18] and in a mouse model of melanoma treated with ACT [26]. Neutrophils infiltrating draining lymph nodes during infection do not enter the parenchyma [32]. Instead, they localize to the medulla and, when activated, enter the interfollicular zone. In the context of early stage HNC, it was found that neutrophil accumulation in T cell rich zones of metastasis-free tdLNs was positively correlated with five-year survival in HNC patients [18]. However, cancer cells reprogrammed the neutrophils toward an immunosuppressive phenotype upon disease progression. How these context-dependent findings translate to neutrophil dynamics and spatial organization in tdLN versus non-tumor-draining lymph node (non-tdLN) during ACT is not yet clear.

Here, we investigate the reactive neutrophil response and the neutrophil–CD8^+^ crosstalk in the lymph nodes during ACT to obtain a more holistic understanding of the cancer-immune interaction. We leverage a murine melanoma model with ACT as previously described [26, 33]. We longitudinally quantify neutrophils and adoptively transferred CD8^+^ T cell subset dynamics over a 14 day period after treatment initiation in lymph nodes from three distinct anatomical regions. Our results suggest that neutrophils have an increased relative abundance and a distinct spatial distribution in the non-tdLN as compared to the tdLN. An increased neutrophil abundance was positively associated with terminally differentiated effector CD8^+^ T cells (T_EFF_ cells), as compared to the longer-lived central memory CD8^+^ T cells (T_CM_ cells). Furthermore, we show that innate immune signaling in the tumor promotes neutrophil responses, and importantly augments differences in neutrophil infiltration between the tdLN and non-tdLN.

## Results

### Neutrophils infiltrate predominantly into the distant contralateral lymph node (clLN) compared to the tdLN during ACT

To investigate the dynamics of different immune cell populations during ACT, C57BL/6 mice were subcutaneously (s.c.) inoculated with a syngeneic melanoma cell line (HC.PmelKO.CDK4R24C-NFhgp100) that was previously established by a method termed CRISPitope (CRISPR-assisted insertion of epitopes) [34, 35] (**Figure 1A**). CRISPitope enables the generation of tumor cells expressing model CD8^+^ T cell epitopes fused to endogenously encoded gene products of choice. In this case, the endogenously expressed oncogenic allele of CDK4 (CDK4R24C) was rendered immunogenic in a neoepitope-like manner by fusing the human gp100 model epitope (hgp100) to its C-terminus. Pmel, the gene encoding for the mouse gp100 was ablated by CRISPR-Cas9 (HC.PmelKO) prior to CRISPitope engineering. Even though the mouse gp100 epitope poorly binds to H2-Db, the respective MHC class I allele in C57BL/6 mice, we considered the knockout of gp100 (Pmel) as preferable because it established a defined genetic background for our ACT protocol. Monoclones from the polyclonal culture of the HC.PmelKO.CDK4R24C-NFhgp100 melanoma cell line [35] were then established and used in the present study. ACT was started when tumor size reached 3–5 mm in diameter. The ACT treatment consisted of a single round of cyclophosphamide (Cy) treatment, followed the next day by intravenous (i.v.) injection of 2 million Pmel-1 T cells, recognizing the hgp100 epitope presented by H2-Db and isolated from spleens of naïve Pmel-1 T-cell receptor (TCR) transgenic mice. Pmel-1 T cell injection was accompanied by *in vivo* activation of the cells by recombinant adenoviral vector Ad-hgp100 as previously described [35–37]. Pmel-1 T cells specifically recognize the hgp100 epitope presented by H2-Db. The treatment was completed with innate immune activation via intratumoral (i.t.) injection of CpG/Polyinosinic:polycytidylic acid (CpG/Poly(I:C)). Cohorts of mice were sampled prior to tumor challenge (Naïve), when tumors reached a diameter of 3-5 mm (Tumor-bearing), one day post-Cy treatment (Cy) and at days 3, 7, and 14 after the start of ACT, as well as upon relapse. Three lymph nodes from distinct anatomical regions were collected from each mouse (**Figure 1B**) and each lymph node was used for either multi-parametric flow cytometry or multiplex immunofluorescence (**Supplementary Table S1**). The lymph nodes considered in this study are: i) the inguinal lymph node right (inLNr), which is the tumor-draining lymph node (tdLN) in this model, ii) the inguinal lymph node left (inLNl), which is the contralateral lymph node (clLN) to the tdLN and thus considered the non-tumor-draining lymph node (non-tdLN), and finally iii) the brachial lymph node right (brLNr), which is the intermediate lymph node (intLN) between the tdLN and the clLN considering its position distant from the tdLN but belonging to the same ipsilateral lymphatic draining basin. Hereafter, will refer to lymph nodes using the functional, drainage-based nomenclature. To determine the time point for the treatment initiation and to monitor mouse well-being, tumor size was measured three times per week.

**Figure 1.**
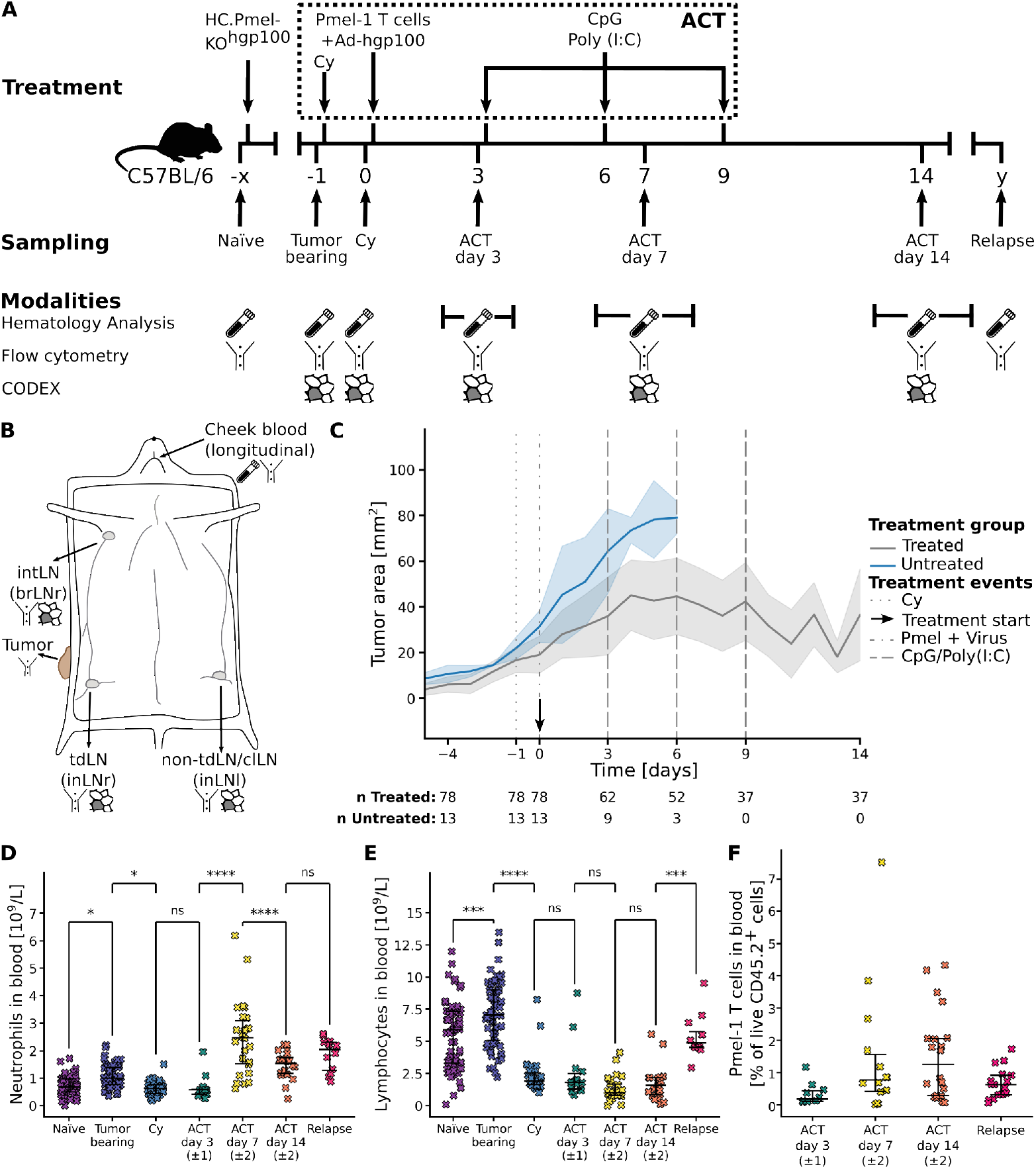
Assessment of tumor establishment and Pmel-1 T-cell engraftment upon ACT. **(A)** Experimental setup and sampling strategy of Pmel-1 ACT in C57BL/6 mice bearing HC.PmelKO.CDK4R24C-NFhgp100 melanomas. Cy: cyclophosphamide treatment. Ad-hgp100: recombinant adenovirus expressing human gp100. CpG/Poly(I:C): CpG/Polyinosinic:polycytidylic acid, innate immune ligands. The complete ACT treatment is indicated within the dashed box. Time points x and y vary between mice. **(B)** Schematic of mouse with lymph nodes and tumor indicated. tdLN: tumor-draining lymph node (inLNr), intLN: intermediate lymph node (brLNr), non-tdLN: non-tumor-draining lymph node (clLN or inLNl). Icons indicate the modalities obtained per tissue as in **(A). (C)** Tumor growth trajectories of all mice. Mice in the treatment group received ACT at indicated time points and were sacrificed at the sampling time points in **(A).** Untreated mice were sacrificed when their tumor reached 8–10 mm in diameter. The shaded area indicates *±*1 standard deviation from the population mean. **(D, E)** White blood cell quantification in cheek blood for neutrophils **(D)** and lymphocytes **(E)** using Mindray BC-5000 Vet Hematology analyzer. **(F)** Pmel-1 T cell concentrations in the blood at sacrifice, measured using flow cytometry. Unpaired tests in **(D-F)** performed using ANOVA with Tukey’s honestly significant difference (HSD) for family-wise error correction. Non-significant associations in **(F)** are not shown.

Our assessment of the tumor size revealed varying dynamics between mice. Around 20% of mice never developed a tumor or had minor tumor growth, but cleared it before it reached the treatment size (**Supplementary Figure S1, Supplementary Table S1**). In the remaining 80%, established tumors responded to ACT and were effectively controlled until day 14 (**Figure 1C**). Assessment of white blood cell populations upon Cy treatment revealed an immediate depletion of peripheral blood neutrophil and lymphocyte concentrations (**Figures 1D, E**). In contrast, the fraction of those cells in the lymph nodes remained mostly stable upon Cy treatment (**Supplementary Figures S2A, B**). ACT was followed by an increase in neutrophil count at day 7 post initiation (**Figure 1D**). (**Figure 1E**). Lymphodepletion in the peripheral blood allowed the Pmel-1 T cells to sustainably engraft in the host, as shown by their presence in the blood (**Figure 1F**) [38].

Next, we assessed changes in the lymph nodes and tumor tissue upon ACT. The Pmel-1 T cells accumulated in the lymph nodes between day 3 and 7 of ACT, preferentially the tdLN (**Supplementary Figure S2C**). The neutrophils also infiltrated the lymph nodes starting from ACT day 3 but mainly between day 7 and 14 (Supplementary **Figure S2A**). Interestingly, a higher fraction of neutrophils infiltrated the clLN (non-tdLN) compared to the tdLN (**Figures 2A-C**). The intermediate lymph node (intLN) showed a similar but slightly less pronounced trend to the clLN (**Supplementary Figure S2A-C**), and will be mostly omitted for brevity. The Pmel-1 T cells in the clLN exhibited predominantly T_EFF_ effector phenotypes (CD44^+^CD62L^-^), whereas in the tdLN the central memory T_CM_ phenotype (CD44^+^CD62L^+^) was predominant (**Figures 2E, F**) [39]. The T_EFF_ population emerged in the tumor mainly at ACT day 7 and remained the predominant Pmel-1 population there throughout the experiment (**Supplementary Figure S2D**). In contrast, only approximately 5% of the Pmel-1 T cells in the tumor were T_CM_ cells, increasing in abundance slightly between ACT day 7 and 14 (**Supplementary Figure S2E**). The T_CM_ phenotype is linked to long-term persistence of transferred T cells [40]. Its predominance among the Pmel-1 phenotypes in the tdLN in our experiments is consistent with this concept.

**Figure 2.**
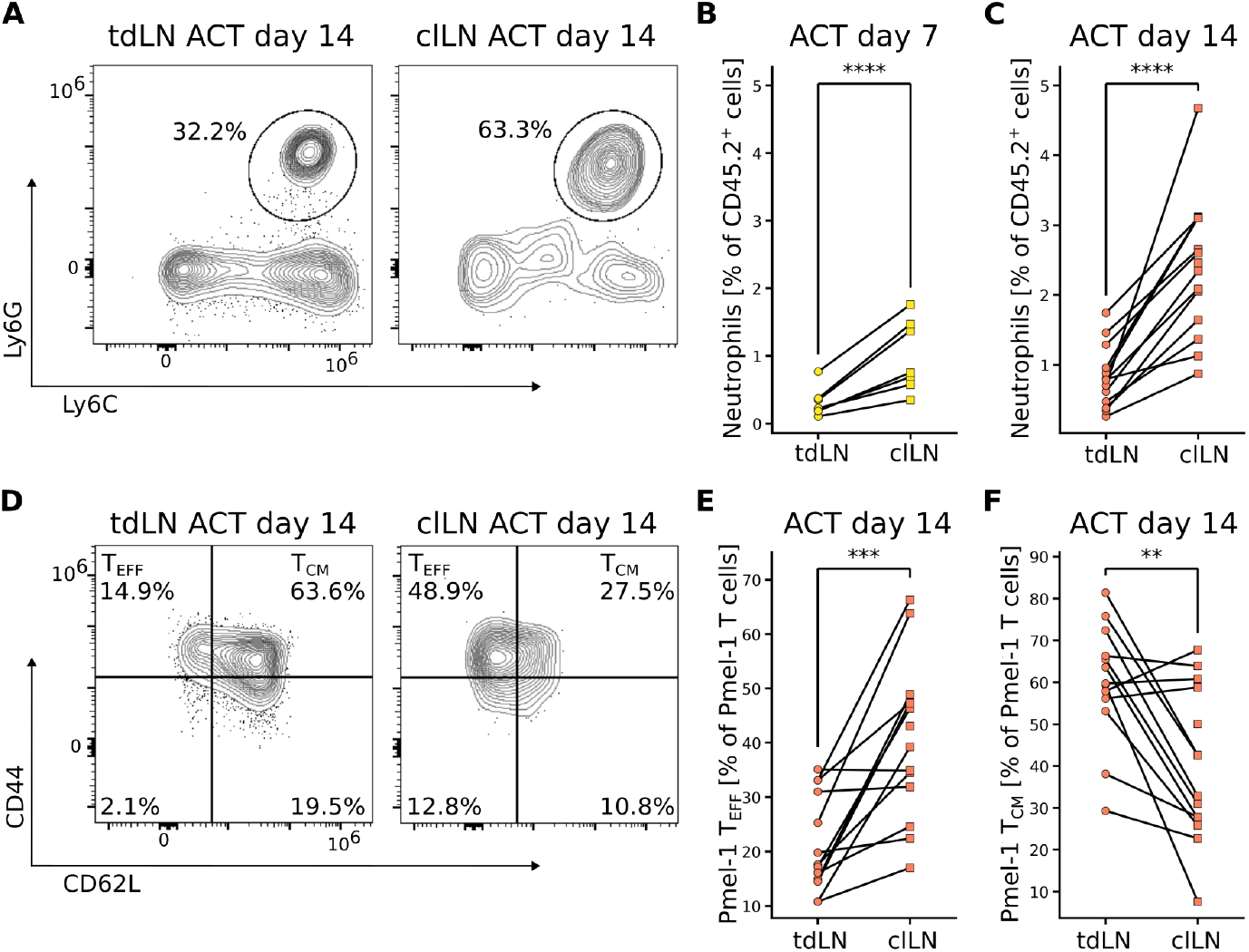
Neutrophil and T cell abundance in the clLN and tdLN. **(A)** Representative flow cytometric plots showing Ly6C against Ly6G expression on CD45.2^+^CD3^-^ cells for neutrophil phenotyping. **(B, C)** Quantification of the relative neutrophil abundance in treatment group ACT day 7 **(B)** and day 14 **(C). (D)** Representative flow cytometric plots showing CD62L against CD44 expression on CD45.2^+^CD8^+^CD90.1^+^ cells for CD8^+^ T cell subtyping. **(E, F)** Quantification of the Pmel-1 T_EFF_ **(E)** and T_CM_ **(F)** subsets in treatment group ACT day 14. Paired tests in **(B, C, E, F)** performed using a ratio-paired T test.

In summary, we observed neutrophils infiltrating both the tdLN and clLN, whereby their fraction was higher in the non-tdLN (clLN). The tdLN contained a higher fraction of beneficial T_CM_ cells.

### Spatial analysis reveals enhanced T cell zone infiltration in the tdLN compared to the clLN

To elucidate the role that the neutrophils play in the lymph nodes upon ACT and to find out how they interact with CD8^+^ T cells, we investigated them in their spatial context during the course of ACT treatment. To this end, we performed multiplex immunofluorescence imaging using the PhenoCycler (formerly co-detection by imaging (CODEX)) platform on the remaining lymph nodes by applying a broad immune cell panel of thirty markers for state and functional phenotyping (**Figures 3A-F, Supplementary Figure S3A**).

**Figure 3.**
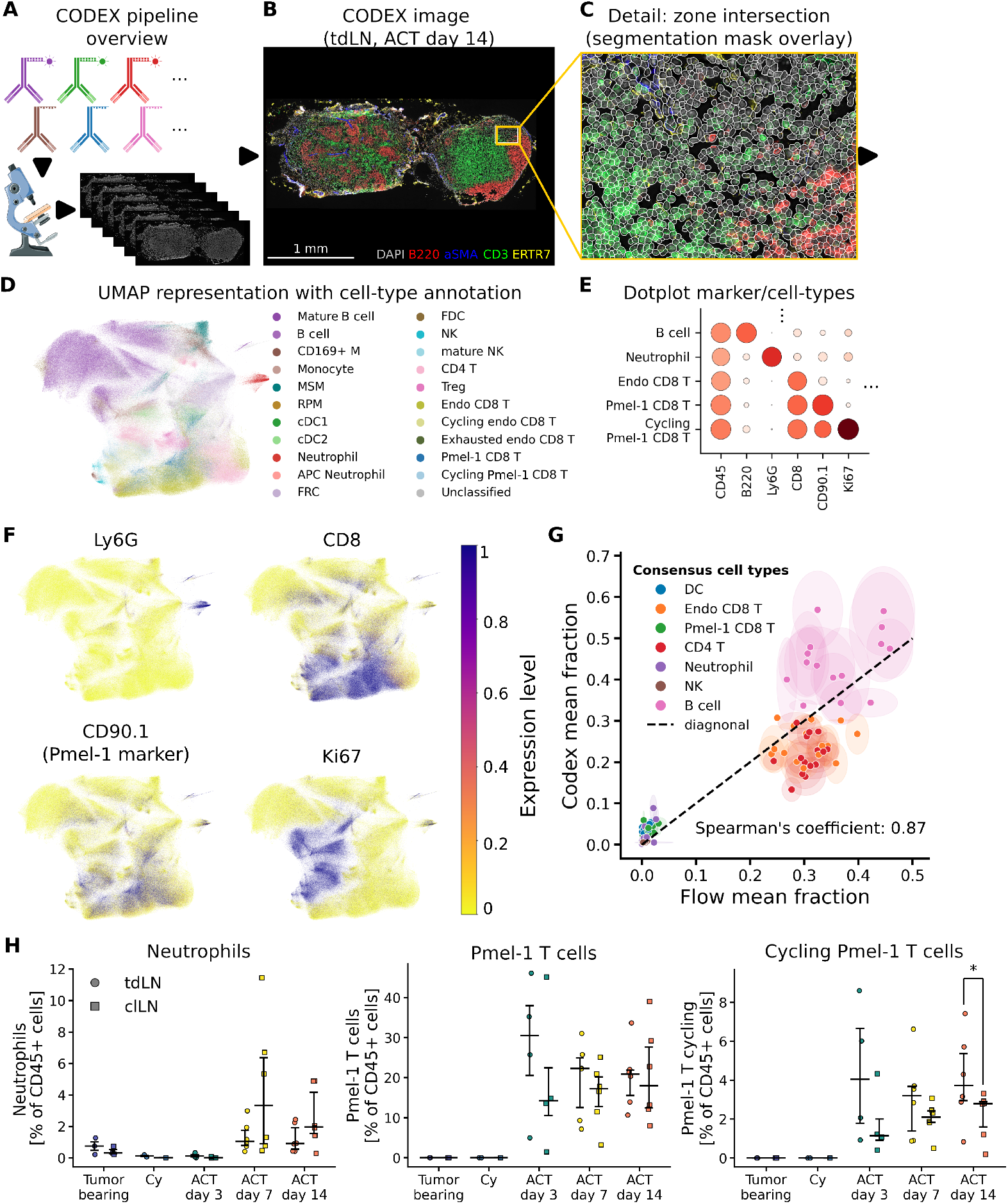
Multiplex immunofluorescence image analysis of lymph nodes. **(A-E)** Overview of the experimental and computational **(B-E)** pipeline. **(B)** tdLN from ACT day 14. Selected markers: DAPI (grey), B220 (red), aSMA (blue), CD3 (green), ERTR7 (yellow). **(C)** Segmentation mask over detail at zone intersection of T cell zone, B cell zone and Medulla of the tdLN in **(B). (D, E)** Sample full UMAP embedding **(D)** and dotplot **(E)** of all multiplex immunofluorescence imaging samples, showing cell type annotation. Dot color in **(E)** is the normalized mean marker expression in the cell type and dot size indicates the fraction of cells expressing the marker in that cell type. **(F)** Expression level of key markers embedded in UMAP space. **(G)** Cross-modality expression alignment. Each dot represents a condition-lymph node combination. **(H)** Quantification of neutrophil and Pmel-1 T cells in the tdLN and clLN per experimental condition in the imaging data. Paired tests in **(H)** performed using a ratio-paired T test. Non-significant associations are not shown.

This analysis identified 22 cell types in their spatial context (**Figure 3D**), with a clear Ki67^+^ CD90.1^+^ (Pmel-1) compartment defining cycling Pmel-1 CD8^+^ T cells (**Figures 3E, F**). Comparison of immune cell populations between the imaging and flow cytometry data revealed a good general alignment, with a Spearman’s rank correlation coefficient of 0.87 (**Figure 3G**). We found that B cells were slightly overrepresented in the imaging data and T cells were slightly underrepresented as compared to the quantification by flow cytometry. In the organ-level distribution of the imaging cell types, the neutrophils and Pmel-1 T cells showed similar trends to their counterparts in the flow cytometry data: Neutrophils appeared more abundant in the clLN and cycling Pmel-1 T cells more in the tdLN (**Figure 3H**).

Beyond the relationship between lymph nodes from distinct anatomical regions, it is important to investigate the spatial distribution of neutrophils within them. To this end, the lymph nodes were classified into three regions based on a spatial graph and cellular neighborhood assignment, followed by a filtering step for increased coherency (**Figure 4A, Supplementary Figure S4**). Comparing a representative tdLN and clLN at day 14 (**Figure 4B**), we observed marked size differences between them. Additionally, in line with the flow cytometry analyses, we observed a higher fraction of neutrophils in the clLN, which appear to be localized mainly in a wide ring outside the T cell zone (**Figures 4C, D**). Exploring this observation further, our quantification across the treatment groups revealed that neutrophil presence was most enhanced in the medulla/subcapsular sinus (SCS)/interfollicular zone (intermediate zone) (**Figure 4E**). In this zone, they comprised six percent of the total cells in the clLN and three percent in the tdLN on average at day 14 after treatment start.

**Figure 4.**
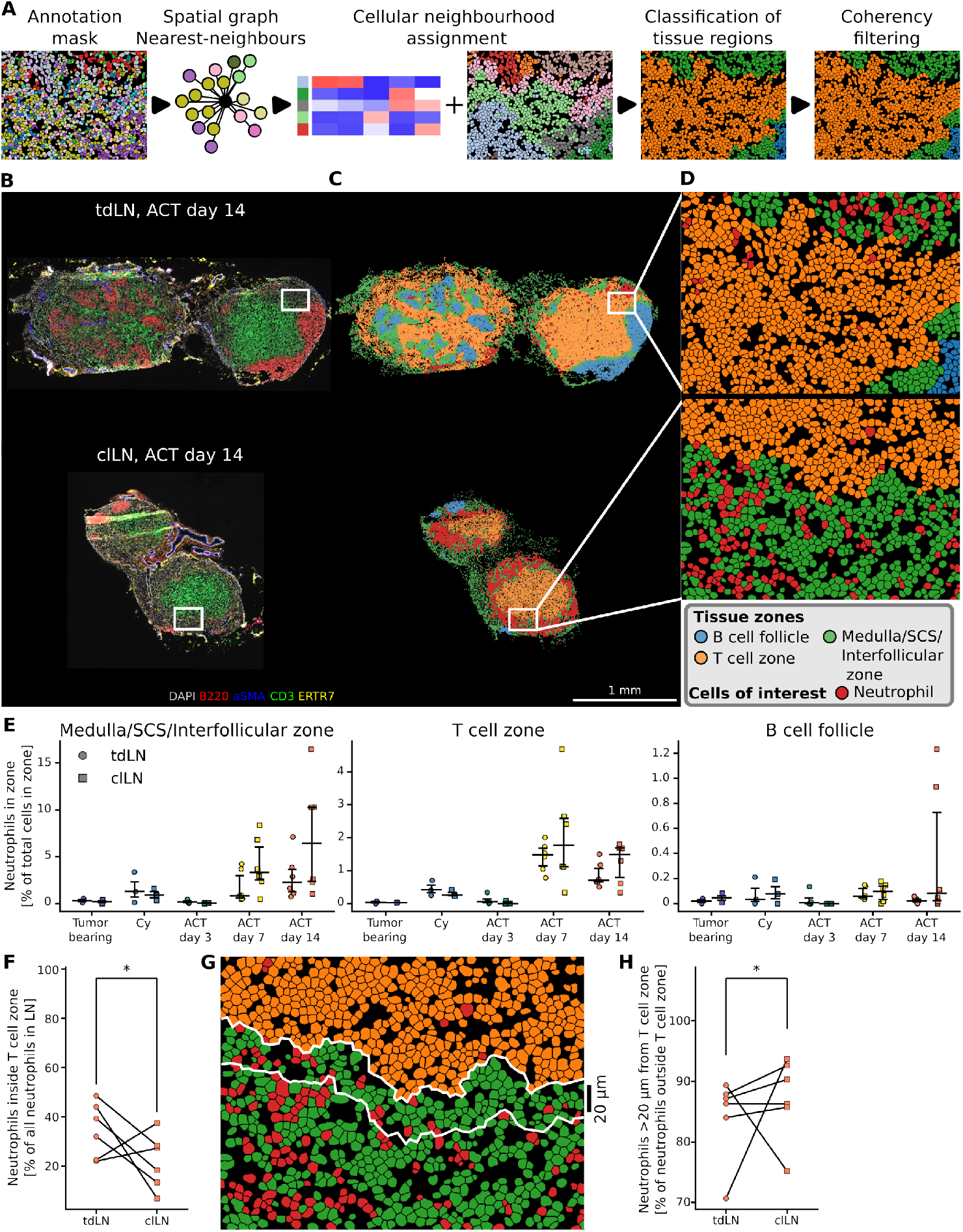
Analysis of neutrophil localization in lymph nodes. **(A)** Overview of spatial analysis workflow. Representative tdLN and clLN at ACT day 14, to scale. Selected markers: DAPI (grey), B220 (red), aSMA (blue), CD3 (green), ERTR7 (yellow). **(C)** Zone classification of the two lymph nodes in **(B).** Neutrophils are indicated in red. **(D)** Zoom of **(C)** with region annotation and neutrophils. **(E)** Neutrophil population in the three tissue zones as a fraction of total cells per zone. **(F)** Neutrophils present in the T cell zone as percentage of all neutrophils, ACT day 14. **(G)** Zoom of clLN from **(D)** with T cell border and 20 µm distance from border indicated in white. **(H)** Neutrophils outside the T cell zone at a 20 µm distance from the border, as a percentage of neutrophils outside the T cell zone. Unpaired tests in **(E)** performed using ANOVA with Tukey’s HSD for family-wise error correction. Non-significant associations in **(E)** are not shown. Paired test for normally distributed data in **(F)** using a ratio-paired T test. Paired test for non-normally distributed data in **(H)** performed using Wilcoxon signed-rank test.

In some mice of the late-stage ACT groups, neutrophils accounted for almost 10% of the total cells in this intermediate zone in the clLN (**Figure 4E**). They also entered the T cell zone, though here they constituted a smaller fraction of this zone. They were hardly present in the B cell follicle. When comparing the lymph node localization at day 14 of ACT, a higher fraction of neutrophils was detected in the T cell zone in the tdLN compared to the clLN (**Figure 4F**). Among the neutrophils that were outside of the T cell zone, those in the clLN seemed to distance themselves from the T cell zone, appearing predominantly more than 20 *µ*m (approximately five cell diameters) away from the T cell zone border (**Figures 4G, H**).

In summary, neutrophils localized mainly outside the T cell zone but were more commonly found there in the tdLN than the clLN. In the clLN, they instead maintained a distance of approximately five cell diameters from the T cell zone.

### Profound changes in lymph node sizes and architecture facilitate immune response in ACT

The tdLN at day 14 of ACT was enlarged to about two times the diameter of the clLN at that same time point (**Figure 4B**). Establishing how lymph node size and growth dynamics correspond to the immune response is crucial for interpreting the response’s functional impact on the organism. To this end, we mapped the growth of the lymph nodes and their anatomical zones over time and between lymph nodes.

Our assessment of bead-corrected live cell counts in flow cytometry revealed that Cy conditioning was associated with a modest, statistically non-significant reduction in lymph node size across all nodes (**Supplementary Figure S5A**). Despite the size decrease, the lymph node architecture did not change visibly under Cy conditioning (**Supplementary Figure S5B**). At later time points the tdLN grew under the effect of ACT and continued tumor presence (**Figures 5A-C**). From the naïve to the relapse time points, the tdLN grew approximately by one order of magnitude from 10^7^ to 10^8^ cells on average (**Supplementary Figure S5A**). The clLN and intLN sizes remained stable during the course of the experiment. The exception to this was the intLN growing significantly between the ACT day 14 and relapse time points. The tdLN contained significantly more cells as compared to the other two lymph nodes from ACT day 7 onward (**Figure 5D**). At relapse, the tdLN contained an order of magnitude more cells than the clLN.

**Figure 5.**
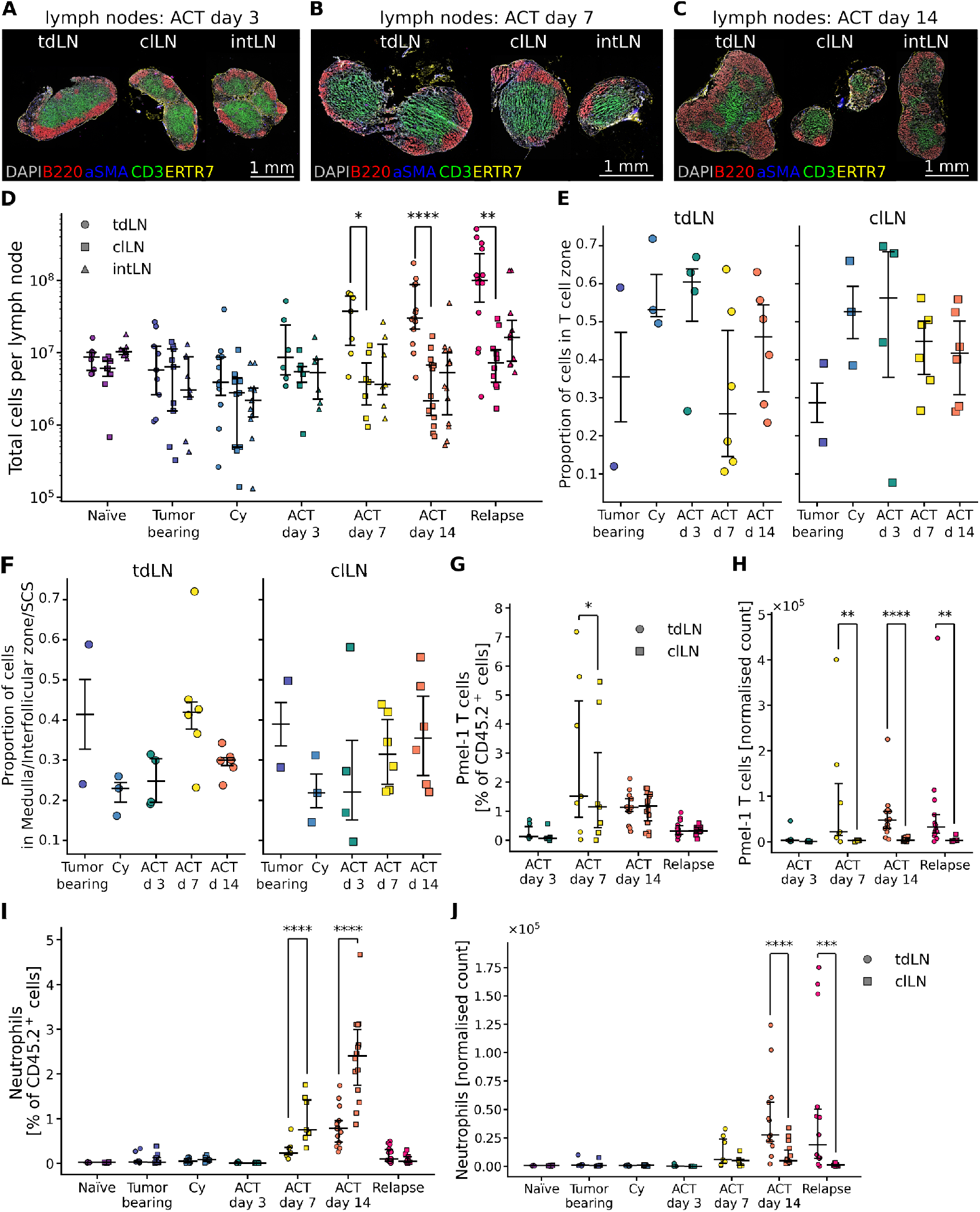
Lymph node size and tissue architecture differences. **(A, B, C)** Lymph node multiplex immunofluorescence imaging images from three representative mice at ACT day 3, 7 and 14. Selected markers: DAPI (grey), B220 (red), aSMA (blue), CD3 (green), ERTR7 (yellow). **(D)** Quantification of lymph node size differences based on total live cell counts in flow cytometry. **(E, F)** Proportion of total cells present in the T cell **(E)** and intermediate zones, as measured by multiplex immunofluorescence imaging. **(G-J)** Pmel-1 T cells **(G, H)** and neutrophils **(I, J)** as a fraction of CD45.2^+^ cells **(G, I)** and as normalized count **(H, J)** in the tdLN and clLN, measured using flow cytometry. Paired tests between tdLN and clLN **(D, G, H, I, J)** performed using a ratio-paired T test. Unpaired tests **(E, F)** performed using ANOVA with Tukey’s HSD for family-wise error correction. Non-significant associations are not shown.

We examined which lymph node anatomical zones contributed to the observed increase in cell number. The proportion of cells that make up the T cell zone increased early after treatment start. At ACT days 7 and 14, this growth was overshadowed by the intermediate zone in both the tdLN and clLN (**Figures 5E, F**). However, variability between mice was large and no significant correlations could be observed. The B cell follicle remained proportionally stable throughout treatment (**Supplementary Figure S5C, D**). These size differences mean that what seems to be a small effect when shown as a percentage could actually obscure a large increase or decrease in an immune cell population, with possible profound biological implications. For example, the fraction of Pmel-1 T cells within the CD45.2^+^ population was similar between the tdLN and the clLN at day 14 and at relapse (**Figure 5G**). In contrast, absolute counts revealed that Pmel-1 T cells were significantly more abundant in the tdLN at ACT days 7 and 14 and at relapse (**Figure 5H**). This underscores the importance of the tdLN as main reservoir of the anti-tumor T cell population. The neutrophils showed an opposite trend: while they were relatively more abundant in the clLN at day 7 and 14 (**Figure 5I**), their absolute counts remained lower than those observed in the rapidly growing tdLN (**Figure 5J**). In the relapse condition, the neutrophil fraction was comparable between the lymph nodes.

To summarize, we observed significant size increases of the tdLN during ACT and emphasized the need to take absolute cell abundances into account when assessing the effect of expanding immune cell populations on the organism.

### Intratumoral innate immune stimulation triggers infiltration of neutrophils in the lymph nodes and promotes T_CM_ cells in the tdLN

We observed previously that fewer Pmel-1 T cells in the clLN exhibited a beneficial T_CM_ phenotype and that the clLN had a higher neutrophil fraction (**Figure 2**). It remained unclear whether this shift was caused by a difference in tumor antigen exposure between these lymph nodes, by a different level of exposure to i.t. CpG/Poly(I:C) innate immune injection, or both. To investigate this, we omitted the CpG/Poly(I:C) stimulation at day 3, 6 and 9 of the ACT protocol and sampled tissues at ACT day 7 and 14 (**Figure 6A**). Importantly, previous work established that i.t. CpG/Poly(I:C) application is in particular critical for the long-term immune surveillance capacity of this combinatorial ACT regimen [36].

**Figure 6.**
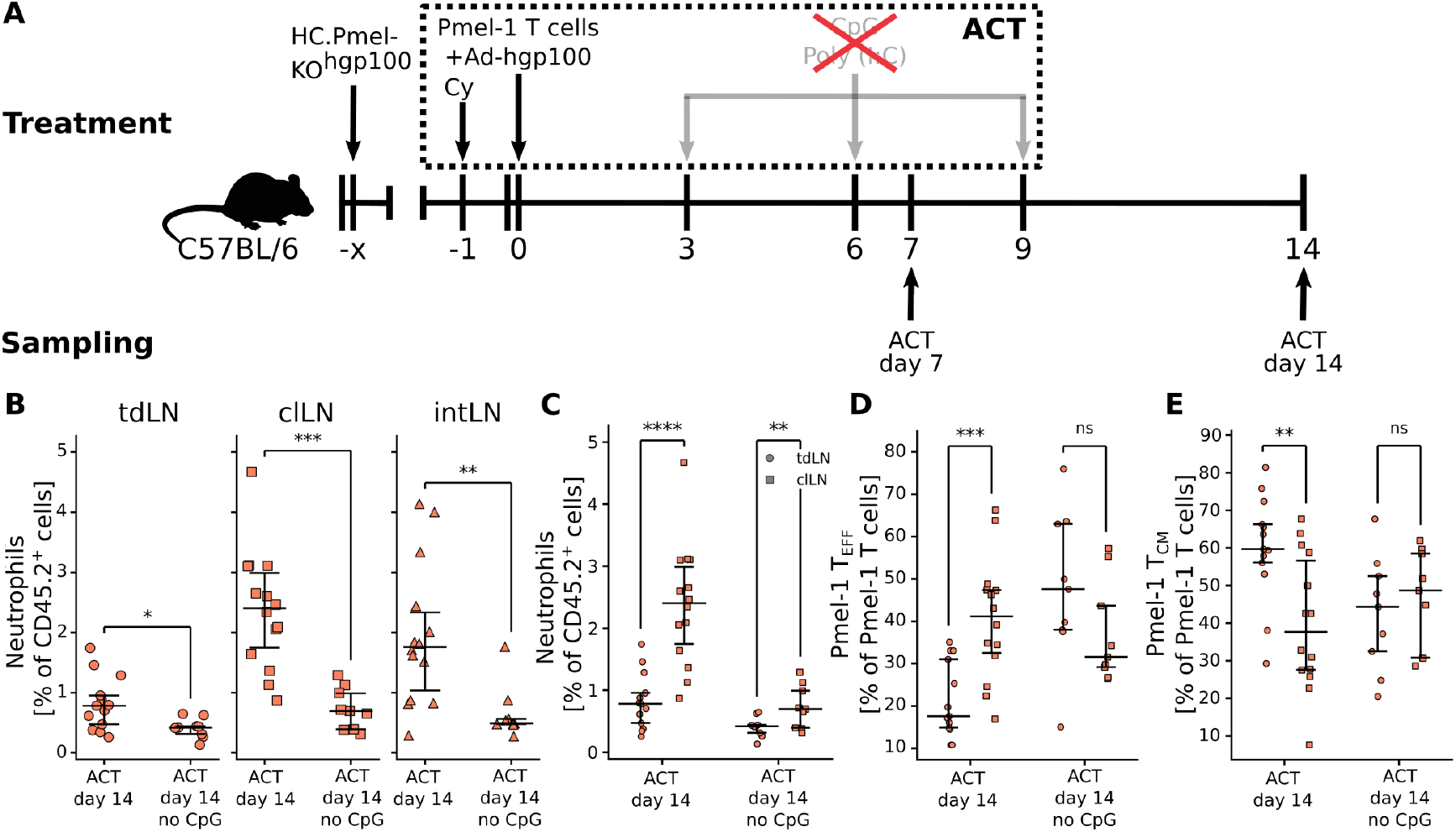
Omission of innate immune stimulation in ACT day 7 and 14. **(A)** Experimental setup and sampling strategy of Pmel-1 ACT without CpG/Poly(I:C) treatment in C57BL/6 mice bearing HC.PmelKO.CDK4R24C-NFhgp100 melanomas. **(B)** Neutrophil abundance with and without CpG/Poly(I:C) treatment at day 14, per lymph node. **(C-E)** Neutrophil, T_EFF_ and T_CM_ concentrations in the lymph nodes at day 14 of ACT with and without innate immune stimulation. Unpaired test in **(B)** performed using ANOVA with Tukey’s HSD for family-wise error correction. Paired tests **(C-E)** performed using a ratio-paired T test. All data was obtained using flow cytometry.

Our assessment of tumor burden in the no-CpG/Poly(I:C) condition revealed similar control of tumor growth as compared conditions with CpG/Poly(I:C) until day 14 (**Supplementary Figures S1J-L**). This is not in disagreement to published work, because the short time window precludes conclusions about long-term therapeutic effects. Lymph node sizes were also similar with and without innate immune stimulation (**Supplementary Figure S6A**). Removal of innate immune stimulation reduced the neutrophil abundance in all lymph nodes (**Figure 6B**). Additionally, the pronounced difference in the neutrophil fraction between the tdLN and the clLN was reduced at ACT day 14 without CpG/Poly(I:C) (**Figure 6C, Supplementary Figure S6B**). Among the Pmel-1 T cells, the T_EFF_ cells showed a similar trend. When omitting innate immune stimulation, at day 14 the difference in T_EFF_ abundance between the tdLN and clLN lymph nodes disappeared (**Figure 6D**). This was mainly due to the significant increase of the T_EFF_ fraction in the tdLN under this condition (**Supplementary Figure S6C**). In addition, the differences in T_CM_ concentration proportions between the clLN and tdLN also disappeared at ACT day 14 without innate immune stimulation (**Figure 6E, Supplementary Figure S6D**). Interestingly, despite these changes in the lymph nodes, the T_EFF_ and T_CM_ populations in the tumor were unchanged (**Supplementary Figures S6E, F**).

To summarize, omitting innate immune stimulation did not change tumor growth dynamics early after treatment start until day 14. It also did not change Pmel-1 T cell populations in the tumor or lymph node sizes. However, without innate immune stimulation, the reactive neutrophil infiltration into all lymph nodes was reduced, which was accompanied by an increase of the T_EFF_ population and a decrease of the T_CM_ population in the tdLN. Thus, our results revealed that i.t. innate immune activation by CpG/Poly(I:C) is important to promote the establishment of the favorable Pmel-1 T_CM_ population in the tdLN, which is known to be critical for long-term persistence.

## Discussion

In the tdLN under ACT, neutrophils acquire an immunosuppressive phenotype [26]. However, a holistic understanding of neutrophil behavior during ACT and the neutrophil-CD8^+^ T cell crosstalk is still incomplete. To address this, we conducted a multi-compartment investigation of lymph nodes from distinct anatomical regions, quantifying neutrophil and Pmel-1 T cells in three different lymph nodes in melanoma-challenged mice treated with ACT. In particular, we focused on the early on-treatment window until day 14 after ACT start to capture the immune cell dynamics in the expansion phase of the adoptively transferred Pmel-1 T cells.

Among the three lymph nodes, we observed an enhanced neutrophil response in the non-tdLN (clLN). This was contrary to our expectation, because we assumed that the tumor-derived cytokine milieu promotes neutrophil infiltration into the tdLN. The enhanced neutrophil presence might be highly relevant, given that previous studies including ours have found that neutrophils in similar circumstances acquire immunosuppressive properties [25, 26, 31, 41, 42]. These immunosuppressive neutrophils can influence functional T cell differentiation either indirectly by promoting CD4^+^ cells to differentiate into effector regulatory T cells [43, 44] or directly, by contact-dependent suppression of CD8^+^ T cell functions [18, 26, 45, 46]. Accordingly, this leads to reduced fitness of the T cell population and an impaired anti-tumor response [22, 25, 27–29].

Following our initial observation of enhanced neutrophil presence in the clLN, we investigated the spatial distribution of the neutrophils within the lymph nodes. We found that the majority of neutrophils were localized outside of the T cell zone, mainly in the medulla. In the tdLN, neutrophils were more likely to be in the T cell zone than their counterparts in the clLN, which also kept a larger relative distance from the T cell zone. Based on their spatial distribution, our data suggests that the neutrophils in the tdLN have more interactions with T cells exerting immunosuppressive functions via PD-L1 and other mechanisms, as shown in our previous work [35]. However, neutrophils in the T cell zones of the tdLN may have both pro – and anti-tumoral roles depending on their phenotype and stage of the disease, as it was previously observed in tdLN samples from HNC patients [18]. The presence of HLA-DR^+^CD80^+^CD86^+^ICAM1^+^PD^-^L1^-^ neutrophils in the T cell zone was associated with favorable disease outcome, whereas PD-L1^+^ neutrophils predicted poor outcome. Another study found distinct neutrophil subsets in human draining and non-draining lymph nodes for patients with oral squamous cell carcinoma [31]. Here, non-draining lymph nodes contained CD16^high^CD62^high^ neutrophils (mature and non-active), whereas tdLN contained more CD16^high^CD62^dim^ (mature and hyper-activated) neutrophils [47], whose abundance correlated with tumor stage [31].

Using the total amount of live cells measured in flow cytometry as a proxy, we calculated the growth of the tdLN during the treatment time course and compared it to the other lymph nodes. We found that the tdLN rapidly increased in size, up to an order of magnitude larger than its counterparts at the relapse time point. Taking LN size and total cell numbers into account is important to interpret cell frequencies properly, as shown for the adoptively transferred Pmel-1 T cells. Their total number was much higher in the tdLN, whereas their frequency remained comparable between the tdLN and non-tdLN.

Lymph node architecture also appeared to be altered, with relatively slower growth of the T cell zone compared to the intermediate zone. An important caveat to this finding is that compositional changes were estimated from 2D tissue slices. While these were cut in the same way each time to be as representative as possible, 2D imaging remains a tenuous modality for this purpose [48, 49]. An example of the uncertainty this introduced becomes apparent when comparing the live cell counts from flow cytometry and cell counts per zone in multiplex immunofluorescence data. In flow cytometry, the total amount of live cells appeared to decrease upon Cy treatment. This was followed by a steady growth, though the size increase was not significant between conditions up to the relapse time point, a kinetic that is very plausible following chemotherapeutic conditioning. By contrast, the cell counts estimated from multiplex immunofluorescence indicated that lymph nodes increased in size upon Cy treatment. However, the sample size was much lower than for flow cytometry and each section may be subject to sample bias. Because we were aware of this weakness in our multiplex immunofluorescence data, we focused on interpreting relative, compositional changes. We only loosely interpreted absolute differences in this modality, as immune cell frequencies were quite robust between flow cytometry and multiplex immunofluorescence.

Next, we investigated the neutrophil response when omitting the intratumoral innate immune stimulation in the tumor and found that the neutrophil response was diminished in the clLN as compared to the tdLN. However, the relative abundance difference did not disappear entirely. This suggests that the strong neutrophil response in the clLN was caused to a large extent by the innate immune stimulation. The T_CM_ population in the tdLN, previously associated with long-term engraftment [8], was decreased when omitting innate stimulation of the melanoma. It remains to be dissected how neutrophils in the tdLN contribute to this T cell phenotype switch, because activation of innate immune signaling in tumors has many favorable effects on antigen processing and presentation that are critical for promoting local and systemic T cell responses.

In summary, we showed that adoptively transferred tumor-reactive CD8+ T cells rapidly expanded after ACT onset, with the highest proportion of a favorable T_CM_ phenotype in the tdLN. Augmenting innate immune signaling in the tumor promoted neutrophil recruitment, most prominently in non-tdLN. However, the presence of neutrophils in the T cell zone was more pronounced in the tdLN, which also harbored the largest absolute pool of transferred CD8^+^ T cells. Collectively, these data reveal distinct early neutrophil kinetics and spatial organization in tdLN versus non-tdLN during ACT, influenced by innate immune signaling in the tumor. Our work, focusing on the lymph node compartment during ACT, underscores the dynamic behavior, complexity and context-dependency of neutrophil responses. We share all relevant resources to facilitate subsequent analysis as well as modeling studies. This study highlights the importance of expanding beyond the conventional tumor-tdLN axis towards considering the lymphatic system as a whole.

## Methods

### Cell lines

The parental HCmel12 melanoma cell line was established from a primary tumor in the HGF-CDK4^R24C^ mouse as previously described [37]. To generate the HC.PmelKO melanoma monoclonal cell line, the Pmel gene was stably knocked-out in the HCmel12 cell line by CRISPR-Cas9 and subsequently re-established as an *ex vivo* cell line to ensure a more homogenous growth *in vivo* [35]. As previously described, the *ex vivo* HC.PmelKO cell line was then modified using the CRISPitope approach to insert CDK4(R24C) protein containing Neon(N) – Flag(F) tag – hgp100 – T2A – Puromycin resistance gene. Monoclones from these cells (B2) and an *ex vivo* derivative of B2 (1923) were established. The B2 or 1923 cell lines were used in these experiments.

All melanoma cell lines were cultured in Roswell Park Memorial Institute 1640 medium containing GlutaMax (Gibco), further supplemented with 10% Fetal Bovine Serum (FBS) (Gibco), 1% non-essential amino acids (Gibco), 1 mM sodium pyruvate (Gibco), 10 mM HEPES (Roth), 55 µM 2-mercaptoethanol (Gibco), 100 IU/mL penicillin G and 100 µg/mL streptomycin sulfate (Gibco) in a humidified incubator at 37 ^*°*^C with 5% CO_2_. The cell lines used were routinely tested for mycoplasma contamination by PCR.

### Animals

Wild-type C57BL/6J (H-2^b^) mice used in this study were purchased from Charles River. TCR-transgenic Pmel-1 (B6.Cg-Thy1a/Cy Tg(TcraTcrb)8Rest/J) mice expressing an αβTCR specific for amino acids 25-33 of human and mouse gp100 presented by H2-D^b^ were bred in-house as previously described [26, 33, 35].

Mice were housed in individually ventilated cages and humidity-controlled under specific pathogen-free conditions in the animal care facility at the University Hospital Bonn. Experiments were performed with 5–13 week old mice. At the beginning of the experiments, age- and sex-matched mice were randomly allocated to different experimental cohorts. All animal experiments were approved by the local government authorities (LAVE, North Rhine-Westphalia, Germany).

### Subcutaneous tumor inoculation

For tumor inoculation, 2 x 10^5^ HC.Pmelko-NF-hgp100.B2 or HC.Pmelko-NF-hgp100.B2.1923 cells in 100 µL phosphate-buffered saline (PBS) were s.c. injected into the shaved flanks of C57BL/6J mice. Tumor sizes were measured with a caliper three times weekly. Tumor area was calculated with the following equation: area = width x length [mm^2^].

### Adoptive T cell therapy

ACT was performed as previously described [35–37]. Briefly, when transplanted tumors reached a mean of 3-5 mm in diameter, mice were injected intraperitoneally (i.p.) with 100 µg of Cy in PBS per gram of mouse. One day post-Cy injection, 2 x 10^6^ naïve Pmel-1 CD8^+^ T cells (in whole splenocytes isolated from Pmel-1 mice) in 200 µL PBS and 5 x 10^8^ plaque-forming units of recombinant adenovirus expressing human gp100 [36, 50] in 100 µl PBS were injected i.v. and i.p., respectively.

At days 3, 6 and 9 post ACT, 50 µg CpG 1826 (MWG biotech) and 50 µg Polyinosinic:polycytidylic acid (Poly(I:C)) (Invivogen) diluted in 100 µL PBS were administered intratumorally.

Mice were euthanized prior to tumor challenge (Naïve), when the tumor reached 3-5 mm or 8-10 mm in diameter (Tumor-bearing or Untreated), one day post-Cy treatment (Cy) and at days 3, 7 or 14 after start of ACT, as well as upon relapse. Additionally, mice were also euthanized when tumors reached 100 mm^2^ or upon signs of illness in adherence with the local ethical regulations.

### Hematological analysis

Blood samples were collected from the submandibular vein of mice maximum twice weekly up until and at endpoint into Lithium-Heparin-coated Microvette 300 µl tubes (Microvette CB300 LH #16.443) and analyzed on Mindray BC-5000 Vet Hematalogy analyzer. The remaining blood sample in the tubes were then used for flow cytometry analysis. Blood samples were obtained at all treatment time points. For ACT day 3 blood samples were taken within a range of *±* 1 days of the time point, for day 7 and 14 within a range of *±* 2 days.

### Flow cytometry analysis

For flow cytometry analysis, the lymph nodes and tumors were dissociated mechanically with 5 mL syringe plungers and digested with 1 mg/mL Collagenase D, 20 µg/mL DNase I and 5% FBS in PBS for 30 mins at 37 ^*°*^C. The digested cell suspensions were then filtered through 70 µM strainers and washed with FACS buffer (PBS containing 2% FBS). After centrifugation, cells were resuspended in PBS in preparation for staining in 96-well round bottom microplates (TPP, #92697).

Immunofluorescent staining of single cell suspensions was performed according to standard protocols. To block non-specific binding of immunoglobulin to the Fc receptors and to exclude dead cells, cells were first incubated with purified anti-CD16/CD32 antibody (Biolegend, #101302) and Live/Dead Blue fixable dye (Thermo Fisher #L23105) for 15 mins at room temperature in the dark. After washing once in FACS buffer, the cells were stained with the following fluorophore-conjugated monoclonal antibodies specific for mouse: CD45.2, CD4, Ly6G, B220, CD62L, CD8a, CD44, CD11c, CD3, CD90.1, XCR1, MHCII, CD11b, NK1.1, CD25 and Ly6C (**Supplementary Figures S3C, D, S7A, B**). Counting beads were added to the samples after the final washing step.

Samples were acquired on Cytek Aurora and analyzed with FlowJo™ software v10 (BD Life Sciences). Corrected cell counts *C*_c_ were calculated from measured counts *C*_m_ by determining the correction factor *F*_c_

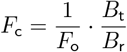

with *F*_o_ the fraction of the organ used in this sample, *B*_*t*_ the total amount of counting beads added and *B*_r_ the amount of beads recovered. For the lymph nodes, the *F*_o_ was 0.5 for the non-ACT cohorts and 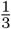 for the ACT cohorts, meaning the full lymph node was divided between the CD8^+^-specific and pan-immune panel or the CD8^+^-specific, pan-immune and CD90.1^+^CD8^+^-specific panels, respectively (**Supplementary Figure S3C, D**). In a few of the larger tdLNs at the relapse time point, a lower fraction was used for flow cytometry and the remaining tissue was frozen. The corrected cell counts were calculated according to *C*_c_ = *C*_m_ · *F*_c_.

### Multiplex immunofluorescence imaging

For the analysis of multiplex immunofluorescence imaging using the PhenoCycler (formerly CODEX) platform, mouse lymph nodes were fixed in BD Cytofix/Cytoperm at 1:4 dilution in PBS for 4 hours at 4 ^*°*^C. Tissues were then washed three times in PBS and dehydrated in 15% sucrose solution in PBS for 1 hour at 4 ^*°*^C. This was followed by further dehydration in 30% sucrose solution in PBS overnight at 4 ^*°*^C. After dehydration, tissues were gently embedded in a cryomold filled with Tissue-Tek OCT (Sakura #4583). Tissues were equilibrated in the Tissue-Tek OCT at room temperature for 20 min and then stored on dry ice for rapid freezing. When completely frozen, tissues were stored at -80 ^*°*^C for long-term storage.

For staining, the dehydrated frozen tissues were cryo-sectioned at 5 µM thickness onto 0.1% poly-L-lysine (Sigma-Aldrich #p8920) coated coverslips. Antibody conjugations and tissue staining were performed as previously described [51]. Briefly, the tissue sections were rehydrated in 1x TBS wash buffer with Tween 20 (Thermo Scientific #28360) and stained using the following antibodies against mouse: Tox, FoxP3, Bcl2, Tbet, CD90.1, CD8, CD103, PD1, CD3, Ly6C, ERTR7, Tcf1, Eomes, Ki67, Ly108, Lag3, CD11c, CD45, CD169, CD69, Vimentin, NKp46, CD31, CD21/35, CD11b, CD4, B220, MHCII, F4/80, αSMA and Ly6G (**Supplementary Figure S3A**).

All antibodies were conjugated in-house to oligonucleotide sequences synthesized by biomers.net GmbH. Images were acquired on a PhenoCycler-Fusion (Akoya Biosciences) coupled to a Zeiss Axio Observer 7 inverted microscope through the CODEX Instrument Manager (CIM; Akoya Biosciences) and ZEN (Zeiss) softwares. Every cycle included the acquisition of DAPI, ATTO 550, DY-647P1 and DY-747P1.

#### Multiplex immunofluorescence image processing

Multiplex immunofluorescence images were processed with an in-house pipeline building on existing methods [52–56].

Following this, initial phenotyping was done based on automatic cell type clustering using a reference phenotyping tree and the mean fluorescence intensity (MFI) per marker as extracted by in-house code. Clustering was done using Leiden clustering [57] using the python package scanpy (v1.9.6) [58, 59]. Clusters were assigned to predefined cell phenotypes. The plausibility of this assignment was evaluated by inspecting the marker MFI per cluster using the `dotplot` and `UMAP` [60] functionality as implemented by scanpy (**Figure 3D, E**).

In cases where the marker intensity profiles of a cluster did not reflect any known phenotype, a subset of the cluster was investigated by inspecting small patches of the image. These patches showed the cell within its direct spatial context, showing the DAPI stain as well as three to five other relevant markers. This visual examination was important to avoid erroneous cell type assignment based on spatial spillover effects or artifacts. After this manual inspection, the cluster was then either assigned to a cell type, discarded as artifact or split further using Leiden clustering. In this way, the dataset was iteratively curated into 22 biologically coherent clusters (**Supplementary Figures S3B**).

#### Spatial analysis

Nearest neighbor graphs were constructed from each image using the Squidpy `spatial_neighbors` function (v1.6.0), with a connection threshold of 99.5% of the total distances and 6 nearest neighbors per point [61]. This left the graph with outliers that were disconnected entirely or only with a few other outliers. Outlying points were then filtered out if they contained less than 1% of the points in the main graph.

For lymph node anatomical zone classification, cells were assigned one of twenty neighborhoods using *k*- nearest neighbor clustering. A nearest neighbor count of *k* = 50 was chosen so that assignment was based on a relatively large spatial context, reflecting tissue anatomy. The twenty clusters were manually assigned to one of three tissue regions based on cell type composition and tissue location. Regions were smoothed by evaluating cell alignment with region labels in neighboring cells (**Figure 4A**). Smoothing was done with a deterministic Metropolis-type algorithm. Briefly, for each cell with one of three tissue region labels, the *k* = 10 nearest neighbors were determined. Among these neighbors, we determined the most common label *L*_most_ and computed

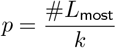

The tissue zone label was deterministically updated according to:

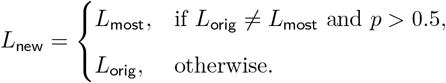

A threshold of 0.5 was chosen in this three-label setting to enable reassignment of surrounded individual points and a slight edge smoothing, but allow label retention in balanced edge regions. Per image, tissue label reassignment was iterated until no changes occurred or the maximum of 10 iterations was exhausted.

Cell type abundance was quantified per tissue region. For region proximity analysis, a concave hull was calculated around the T cell zones and distance of cells to the hull border calculated using the SPACEc `patch_proximity_analysis` function (v0.1.2) using the `border_cell_radius` method [62]. A distance of approximately five cell diameters (20 µm) was chosen to reflect the distance at which cells were no longer able to directly interact with the T cell zone. This distance is larger than what appears to be the plausible direct interaction radius in 2D. However, recent studies have found that interaction distances are overestimated in 2D compared to the underlying 3D tissue structure in human skin and TME immune contexts [63, 64]. Accordingly, a 20 µm distance was chosen to compensate for distances to possible interaction partners potentially being inflated as compared to the actual 3D context. Cells in and outside of this direct interaction zone were counted.

### Flow cytometry-multiplex immunofluorescence imaging comparison

Consensus cell types were defined as cell types present in both modalities. These were grouped by organ and experimental condition. The mean and one standard deviation of the per-sample cell type fractions were determined. Spearman’s rank correlation coefficient was used to determine alignment along the diagonal for these fractions. This nonparametric measure was chosen because it is scale-independent.

### Statistical analysis

Statistical analysis between treatment groups was performed using ANOVA with Tukey’s HSD for family-wise error correction. All data used in these analyses was normally distributed, as tested using the Shapiro-Wilk test. Comparisons within groups between different organs were done using ratio-paired T tests (for normally distributed data) or the Wilcoxon signed-rank test (for non-normally distributed data). Normality was tested using the Shapiro-Wilk test. P-values lower than 0.05 were considered statistically significant. Analysis was done in Python (v3.10) using scipy (v1.15.1), statsmodels (v0.14.4) and scikit posthocs (v0.11.2).

## Data availability

Data are available in Zenodo at https://doi.org/10.5281/zenodo.18712088

## Code availability

The original code for this manuscript is currently attached as supplement and will be uploaded to GitHub and Zenodo after approval.

## Acknowledgments

This work was supported by the Deutsche Forschungsgemeinschaft (DFG, German Research Foundation) under Germany’s Excellence Strategy (EXC 2047—390685813, EXC 2151—390873048), the Horizon Europe - ERC Consolidator Grant 2023 (grant agreement no. 101126146), the German Federal Ministry for Research, Technology and Space (BMFTR) via the project interpretTME (grant agreement no. 031L0308A & 031L0308B), and by the University of Bonn through the Center for Mathematical Life Sciences and via the Schlegel Professorship of J.H.; and by the Deutsche Krebshilfe (German Cancer Aid) project grant 70114292 (Excellence Program) and Deutsche Forschungsgemeinschaft (DFG, German Research Foundation) under Germany’s Excellence Strategy (EXC 2151—390873048) to M.H.

The authors gratefully acknowledge the access to the Marvin HPC cluster of the University of Bonn.

## Author contributions

GV: Data curation, Formal analysis, Methodology (computational), Software, Visualization, Writing – original draft, Writing – review & editing. ME: Data curation, Formal analysis, Methodology (experimental), Investigation, Project administration, Writing – review & editing. MY: Data curation, Investigation, Writing – original draft, Writing – review & editing. LK: Software. RT: Methodology (experimental), Writing – review & editing. SL: Methodology (experimental). SN: Methodology (experimental). DC: Methodology (experimental), Writing – review & editing. TB: Resources. NG: Resources, Writing – review & editing. KT: Funding acquisition, Supervision, Writing – review & editing. JH: Funding acquisition, Supervision, Methodology (computational), Writing – review & editing. MH: Conceptualization, Funding acquisition, Methodology (experimental), Project administration, Supervision, Writing – review & editing. All authors have approved the final manuscript.

## Ethics statement

All animal experiments were approved by the local government authorities (LAVE, North Rhine-Westphalia, Germany) and conducted in accordance with legislation of the national and institutional animal welfare bodies.

## Supplementary material

The Supplementary Material for this article can be found online at: …

## References

1. Albarrán, V. et al. Adoptive T cell therapy for solid tumors: current landscape and future challenges. Frontiers in Immunology 15. https://www.frontiersin.org/journals/immunology/articles/10.3389/fimmu.2024.1352805/full (2024).

2. Rosenberg, S. A. & Dudley, M. E. Adoptive cell therapy for the treatment of patients with metastatic melanoma. Current Opinion in Immunology 21, 233–240. https://www.sciencedirect.com/science/article/pii/S0952791509000259 (2009).

3. Dafni, U et al. Efficacy of adoptive therapy with tumor-infiltrating lymphocytes and recombinant interleukin-2 in advanced cutaneous melanoma: a systematic review and meta-analysis. Annals of Oncology 30, 1902–1913. https://www.sciencedirect.com/science/article/pii/S0923753420325539 (2019).

4. Rohaan, M. W. et al. Tumor-Infiltrating Lymphocyte Therapy or Ipilimumab in Advanced Melanoma. The New England Journal of Medicine 387, 2113–2125. https://www.nejm.org/doi/full/10.1056/NEJMoa2210233 (2022).

5. Li, F. et al. The association between CD8+ tumor-infiltrating lymphocytes and the clinical outcome of cancer immunotherapy: A systematic review and meta-analysis. eClinicalMedicine 41, 101134. https://www.sciencedirect.com/science/article/pii/S2589537021004144 (2021).

6. Krishna, S. et al. Stem-like CD8 T cells mediate response of adoptive cell immunotherapy against human cancer. Science 370, 1328–1334. https://www.science.org/doi/10.1126/science.abb9847 (2020).

7. Gebhardt, T., Park, S. L. & Parish, I. A. Stem-like exhausted and memory CD8+ T cells in cancer. Nature Reviews Cancer 23, 780–798. https://www.nature.com/articles/s41568-023-00615-0 (2023).

8. Klebanoff, C. A. et al. Central memory self/tumor-reactive CD8+ T cells confer superior antitumor immunity compared with effector memory T cells. Proceedings of the National Academy of Sciences 102, 9571–9576. https://www.pnas.org/doi/abs/10.1073/pnas.0503726102 (2005).

9. Delclaux, I. et al. The tumor-draining lymph node as a reservoir for systemic immune surveillance. Trends in Cancer 10, 28–37. https://www.sciencedirect.com/science/article/pii/S2405803323001905 (2024).

10. Prokhnevska, N. et al. CD8+ T cell activation in cancer comprises an initial activation phase in lymph nodes followed by effector differentiation within the tumor. Immunity 56, 107–124.e5. https://www.cell.com/immunity/fulltext/S1074-7613(22)00606-9 (2023).

11. Ganti, S. N. et al. Regulatory B cells preferentially accumulate in tumor-draining lymph nodes and promote tumor growth. Scientific Reports 5, 12255. https://www.nature.com/articles/srep12255 (2015).

12. Fransen, M. F. et al. Tumor-draining lymph nodes are pivotal in PD-1/PD-L1 checkpoint therapy. JCI Insight 3. https://insight.jci.org/articles/view/124507 (2018).

13. Rotman, J. et al. Unlocking the therapeutic potential of primary tumor-draining lymph nodes. Cancer Immunology, Immunotherapy 68, 1681–1688. 10.1007/s00262-019-02330-y (2019).

14. Wei, J. et al. Immune microenvironment of tumor-draining lymph nodes: insights for immunotherapy. Frontiers in Immunology 16. https://www.frontiersin.org/journals/immunology/articles/10.3389/fimmu.2025.1562797/full (2025).

15. Habenicht, L. M. et al. Distinct mechanisms of B and T lymphocyte accumulation generate tumor-draining lymph node hypertrophy. OncoImmunology 5, e1204505. 10.1080/2162402X.2016.1204505 (2016).

16. Schwartz, L. H. et al. Evaluation of lymph nodes with RECIST 1.1. European Journal of Cancer 45, 261–267. https://www.sciencedirect.com/science/article/pii/S0959804908008757 (2009).

17. Harrell, M. I., Iritani, B. M. & Ruddell, A. Tumor-Induced Sentinel Lymph Node Lymphangiogenesis and Increased Lymph Flow Precede Melanoma Metastasis. The American Journal of Pathology 170, 774–786. https://www.sciencedirect.com/science/article/pii/S000294401060898X (2007).

18. Pylaeva, E. et al. During early stages of cancer, neutrophils initiate anti-tumor immune responses in tumor-draining lymph nodes. Cell Reports 40, 111171. https://www.sciencedirect.com/science/article/pii/S2211124722009846 (2022).

19. Xiong, S., Dong, L. & Cheng, L. Neutrophils in cancer carcinogenesis and metastasis. Journal of Hematology & Oncology 14, 173. 10.1186/s13045-021-01187-y (2021).

20. Rosales, C. Neutrophils at the crossroads of innate and adaptive immunity. Journal of Leukocyte Biology 108, 377–396. 10.1002/JLB.4MIR0220-574RR (2020).

21. Cai, W. et al. Neutrophils in cancer: At the crucial crossroads of anti-tumor and pro-tumor. Cancer Communications 45, 888–913. https://onlinelibrary.wiley.com/doi/abs/10.1002/cac2.70027 (2025).

22. Si, Y. et al. Multidimensional imaging provides evidence for down-regulation of T cell effector function by MDSC in human cancer tissue. Science Immunology 4, eaaw9159. https://www.science.org/doi/10.1126/sciimmunol.aaw9159 (2019).

23. Raskov, H. et al. Neutrophils and polymorphonuclear myeloid-derived suppressor cells: an emerging battleground in cancer therapy. Oncogenesis 11, 22. https://www.nature.com/articles/s41389-022-00398-3 (2022).

24. Law, A. M. K., Valdes-Mora, F. & Gallego-Ortega, D. Myeloid-Derived Suppressor Cells as a Therapeutic Target for Cancer. Cells 9, 561. https://www.mdpi.com/2073-4409/9/3/561 (2020).

25. Jaillon, S. et al. Neutrophil diversity and plasticity in tumour progression and therapy. Nature Reviews Cancer 20, 485–503. https://www.nature.com/articles/s41568-020-0281-y (2020).

26. Glodde, N. et al. Reactive Neutrophil Responses Dependent on the Receptor Tyrosine Kinase c-MET Limit Cancer Immunotherapy. Immunity 47, 789–802.e9. https://www.sciencedirect.com/science/article/pii/S1074761317304235 (2017).

27. Kaltenmeier, C. et al. Neutrophil Extracellular Traps Promote T Cell Exhaustion in the Tumor Microenvironment. Frontiers in Immunology 12. https://www.frontiersin.org/journals/immunology/articles/10.3389/fimmu.2021.785222/full (2021).

28. Emmons, T. R. et al. Mechanisms Driving Neutrophil-Induced T-cell Immunoparalysis in Ovarian Cancer. Cancer Immunology Research 9, 790–810. 10.1158/2326-6066.CIR-20-0922 (2021).

29. Tie, Y. et al. Immunosuppressive cells in cancer: mechanisms and potential therapeutic targets. Journal of Hematology & Oncology 15, 61. 10.1186/s13045-022-01282-8 (2022).

30. Lok, L. S. C. et al. Phenotypically distinct neutrophils patrol uninfected human and mouse lymph nodes. Proceedings of the National Academy of Sciences of the United States of America 116, 19083–19089. https://pmc.ncbi.nlm.nih.gov/articles/PMC6754587/ (2019).

31. Ekstedt, S. et al. Phenotypical differences of neutrophils patrolling tumour-draining lymph nodes in head and neck cancer. British Journal of Cancer 131, 1893–1900. https://www.nature.com/articles/s41416-024-02891-5 (2024).

32. De Castro Pinho, J. & Förster, R. Lymph-Derived Neutrophils Primarily Locate to the Subcapsular and Medullary Sinuses in Resting and Inflamed Lymph Nodes. Cells 10, 1486. https://www.mdpi.com/2073-4409/10/6/1486 (2021).

33. Landsberg, J. et al. Melanomas resist T-cell therapy through inflammation-induced reversible dedifferentiation. Nature 490, 412–416. https://www.nature.com/articles/nature11538 (2012).

34. Effern, M. et al. CRISPitope: A generic platform to model target antigens for adoptive T cell transfer therapy in mouse tumor models. STAR Protocols 3, 101038. https://europepmc.org/article/pmc/8755566 (2022).

35. Effern, M. et al. Adoptive T Cell Therapy Targeting Different Gene Products Reveals Diverse and Context-Dependent Immune Evasion in Melanoma. Immunity 53, 564–580.e9. https://www.sciencedirect.com/science/article/pii/S1074761320303149 (2020).

36. Kohlmeyer, J. et al. Complete regression of advanced primary and metastatic mouse melanomas following combination chemoimmunotherapy. Cancer Research 69, 6265–6274. https://aacrjournals.org/cancerres/article/69/15/6265/549845/Complete-Regression-of-Advanced-Primary-and (2009).

37. Bald, T. et al. Ultraviolet-radiation-induced inflammation promotes angiotropism and metastasis in melanoma. Nature 507, 109–113. https://www.nature.com/articles/nature13111 (2014).

38. Lickefett, B. et al. Lymphodepletion – an essential but undervalued part of the chimeric antigen receptor T-cell therapy cycle. Frontiers in Immunology 14, 1303935. https://pmc.ncbi.nlm.nih.gov/articles/PMC10770848/ (2023).

39. Hamann, D. et al. Phenotypic and Functional Separation of Memory and Effector Human CD8+ T Cells. Journal of Experimental Medicine 186, 1407–1418. 10.1084/jem.186.9.1407 (1997).

40. Busch, D. H. et al. Role of memory T cell subsets for adoptive immunotherapy. Seminars in Immunology 28, 28–34. https://www.sciencedirect.com/science/article/pii/S104453231600004X (2016).

41. Rapoport, B. L. et al. Role of the Neutrophil in the Pathogenesis of Advanced Cancer and Impaired Responsiveness to Therapy. Molecules 25, 1618. https://www.mdpi.com/1420-3049/25/7/1618 (2020).

42. Raftopoulou, S. et al. Tumor-Mediated Neutrophil Polarization and Therapeutic Implications. International Journal of Molecular Sciences 23, 3218. https://www.mdpi.com/1422-0067/23/6/3218 (2022).

43. Haist, M. et al. The Functional Crosstalk between Myeloid-Derived Suppressor Cells and Regulatory T Cells within the Immunosuppressive Tumor Microenvironment. Cancers 13, 210. https://www.mdpi.com/2072-6694/13/2/210 (2021).

44. Wang, C. et al. Immunosuppressive JAG2+ tumor-associated neutrophils hamper PD-1 blockade response in ovarian cancer by mediating the differentiation of effector regulatory T cells. Cancer Communications 45, 747–773. https://onlinelibrary.wiley.com/doi/abs/10.1002/cac2.70021 (2025).

45. Michaeli, J. et al. Tumor-associated neutrophils induce apoptosis of non-activated CD8 T-cells in a TNFα and NO-dependent mechanism, promoting a tumor-supportive environment. OncoImmunology 6, e1356965. 10.1080/2162402X.2017.1356965 (2017).

46. Kwantwi, L. B. et al. Tumor-associated neutrophils activated by tumor-derived CCL20 (C-C motif chemokine ligand 20) promote T cell immunosuppression via programmed death-ligand 1 (PD-L1) in breast cancer. Bioengineered 12, 6996–7006. 10.1080/21655979.2021.1977102 (2021).

47. Pillay, J. et al. Functional heterogeneity and differential priming of circulating neutrophils in human experimental endotoxemia. Journal of Leukocyte Biology 88, 211–220. 10.1189/jlb.1209793 (2010).

48. Uhlén, P. & Tanaka, N. Improved Pathological Examination of Tumors with 3D Light-Sheet Microscopy. Trends in Cancer 4, 337–341. https://www.sciencedirect.com/science/article/pii/S2405803318300608 (2018).

49. Ellingson, B. M. et al. Volumetric measurements are preferred in the evaluation of mutant IDH inhibition in non-enhancing diffuse gliomas: Evidence from a phase I trial of ivosidenib. Neuro-Oncology 24, 770– 778 (2022).

50. Jonuleit, H. et al. Efficient transduction of mature CD83+ dendritic cells using recombinant adenovirus suppressed T cell stimulatory capacity. Gene Therapy 7, 249–254 (2000).

51. Black, S. et al. CODEX multiplexed tissue imaging with DNA-conjugated antibodies. Nature Protocols 16, 3802–3835 (2021).

52. Czech, E. et al. Cytokit: a single-cell analysis toolkit for high dimensional fluorescent microscopy imaging. BMC Bioinformatics 20, 448. 10.1186/s12859-019-3055-3 (2019).

53. Forster, B. et al. Complex wavelets for extended depth-of-field: A new method for the fusion of multichannel microscopy images. Microscopy Research and Technique 65, 33–42. https://onlinelibrary.wiley.com/doi/abs/10.1002/jemt.20092 (2004).

54. Peng, T. et al. A BaSiC tool for background and shading correction of optical microscopy images. Nature Communications 8, 14836. https://www.nature.com/articles/ncomms14836 (2017).

55. Muhlich, J. L. et al. Stitching and registering highly multiplexed whole-slide images of tissues and tumors using ASHLAR. Bioinformatics 38, 4613–4621. 10.1093/bioinformatics/btac544 (2022).

56. Stringer, C. & Pachitariu, M. Cellpose3: one-click image restoration for improved cellular segmentation. Nature Methods 22, 592–599. https://www.nature.com/articles/s41592-025-02595-5 (2025).

57. Traag, V. A., Waltman, L. & van Eck, N.J. /From Louvain to Leiden: guaranteeing well-connected communities. Scientific Reports 9, 5233. https://www.nature.com/articles/s41598-019-41695-z (2019).

58. Wolf, F. A., Angerer, P. & Theis, F. J. SCANPY: large-scale single-cell gene expression data analysis. Genome Biology 19, 15. 10.1186/s13059-017-1382-0 (2018).

59. Virshup, I. et al. The scverse project provides a computational ecosystem for single-cell omics data analysis. Nature Biotechnology 41, 604–606. https://www.nature.com/articles/s41587-023-01733-8 (2023).

60. McInnes, L., Healy, J. & Melville, J. UMAP: Uniform Manifold Approximation and Projection for Dimension Reduction 2020. arXiv: 1802.03426[stat]. http://arxiv.org/abs/1802.03426.

61. Palla, G. et al. Squidpy: a scalable framework for spatial omics analysis. Nature Methods 19. Number: 2, 171–178. https://www.nature.com/articles/s41592-021-01358-2 (2022).

62. Tan, Y. et al. SPACEc: a streamlined, interactive Python workflow for multiplexed image processing and analysis. Nature Communications 16, 10652. https://www.nature.com/articles/s41467-025-65658-3 (2025).

63. Ghose, S. et al. 3D reconstruction of skin and spatial mapping of immune cell density, vascular distance and effects of sun exposure and aging. Communications Biology 6, 718. https://www.nature.com/articles/s42003-023-04991-z (2023).

64. Pentimalli, T. M. et al. Combining spatial transcriptomics and ECM imaging in 3D for mapping cellular interactions in the tumor microenvironment. Cell Systems 16, 101261. https://www.sciencedirect.com/science/article/pii/S2405471225000948 (2025).

