## Supplementary Materials for "Longitudinal spatial profiling of neutrophils during adoptive T cell therapy in murine melanoma reveals distinct lymph node infiltration patterns across anatomical sites"

Table S1: Dataset overview. CODEX: co-detection by imaging, FCM: flow cytometry

| **Condition** | **Mice: total** | **Mice: tumor growth** | **Mice: FCM** | **Mice: CODEX** | **FCM: Mean live cell count per LN (±sd)** $[10^6]$ | **CODEX: Mean live cell count per LN slice (±sd)** $[10^3]$ |
|---|---|---|---|---|---|---|
| Naïve | 24 | 0 | 8 | 0 | 7.04 (±3.06) | – |
| Tumor-bearing | 24 | 19 | 9 | 2 | 5.84 (±5.67) | 17.84 (±17.60) |
| Cy | 23 | 16 | 11 | 3 | 3.62 (±5.91) | 30.16 (±18.24) |
| ACT day 3 | 14 | 10 | 6 | 4 | 7.30 (±10.09) | 30.52 (±13.43) |
| ACT day 7 | 15 | 15 | 7 | 6 | 14.07 (±18.14) | 30.93 (±15.73) |
| ACT day 14 | 22 | 20 | 14 | 6 | 17.89 (±29.77) | 33.59 (±18.86) |
| Relapse | 24 | 17 | 14 | 0 | 65.22 (±100.89) | – |
| Untreated | 17 | 13 | 0 | 0 | – | – |
| ACT day 7 no CpG | 6 | 5 | 5 | 0 | 3.83 (±4.06) | – |
| ACT day 14 no CpG | 9 | 9 | 9 | 0 | 19.40 (±29.06) | – |
| **Total** | 178 | 124 | 83 | 21 | 21.92 (±55.19) | 30.46 (±16.75) |

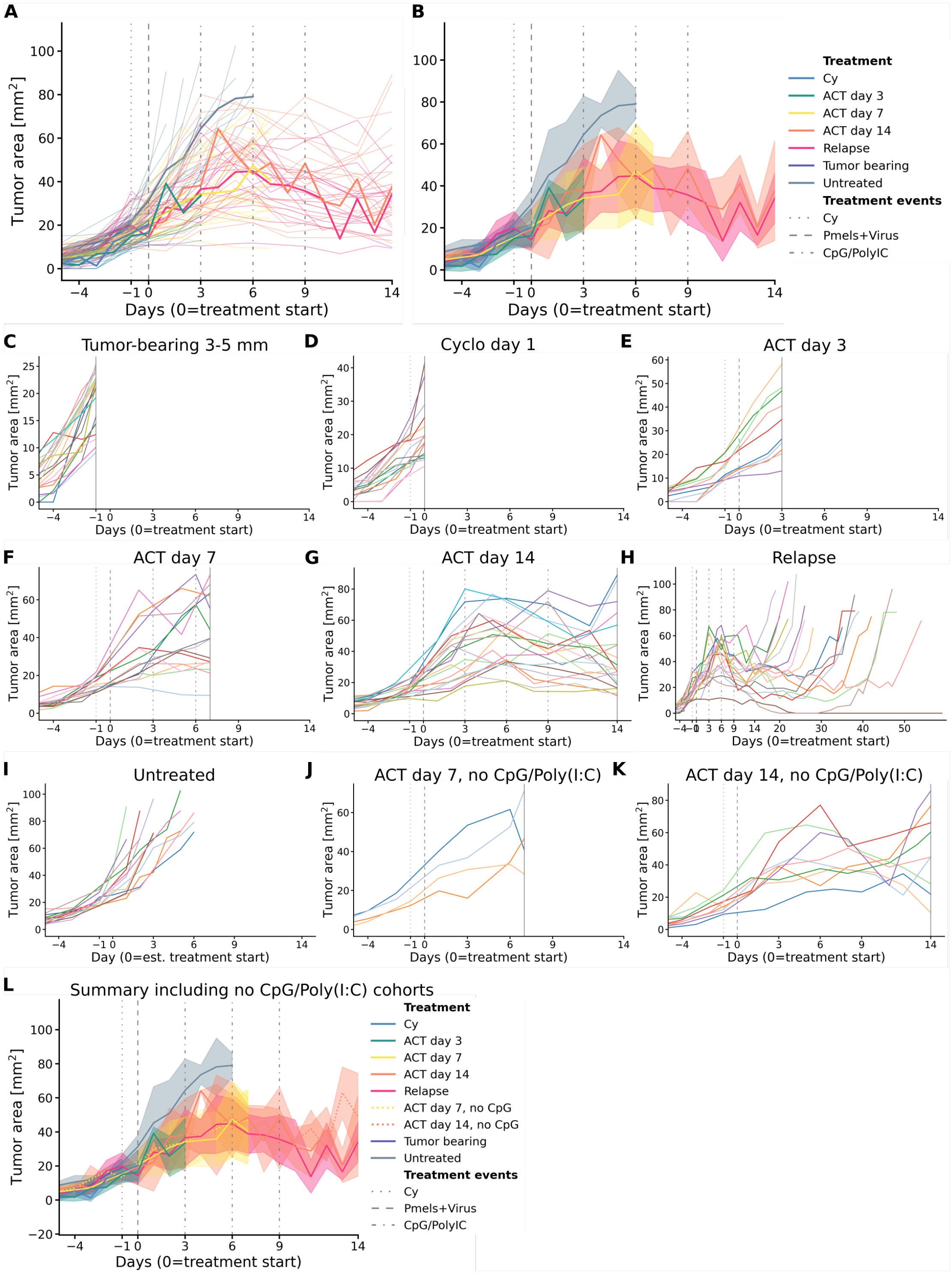


Figure S1: **Tumor growth curves normalized by treatment starting date.** **(A, B)** Individual and averaged trajectories of all non-perturbation conditions. The shaded area in **(B)** indicates ±1 standard deviation from the population mean. **(C-K)** Individual growth trajectories per mouse per condition. The solid line indicates the sacrifice event. **(I)** Treatment start estimated as the day after tumor reached 16 $mm^2$, the mean intended size. **(L)** Averaged trajectories including the conditions without CpG/Polyinosinic:polycytidylic acid (CpG/Poly(I:C)) stimulation. The shaded area indicates ±1 standard deviation from the population mean.

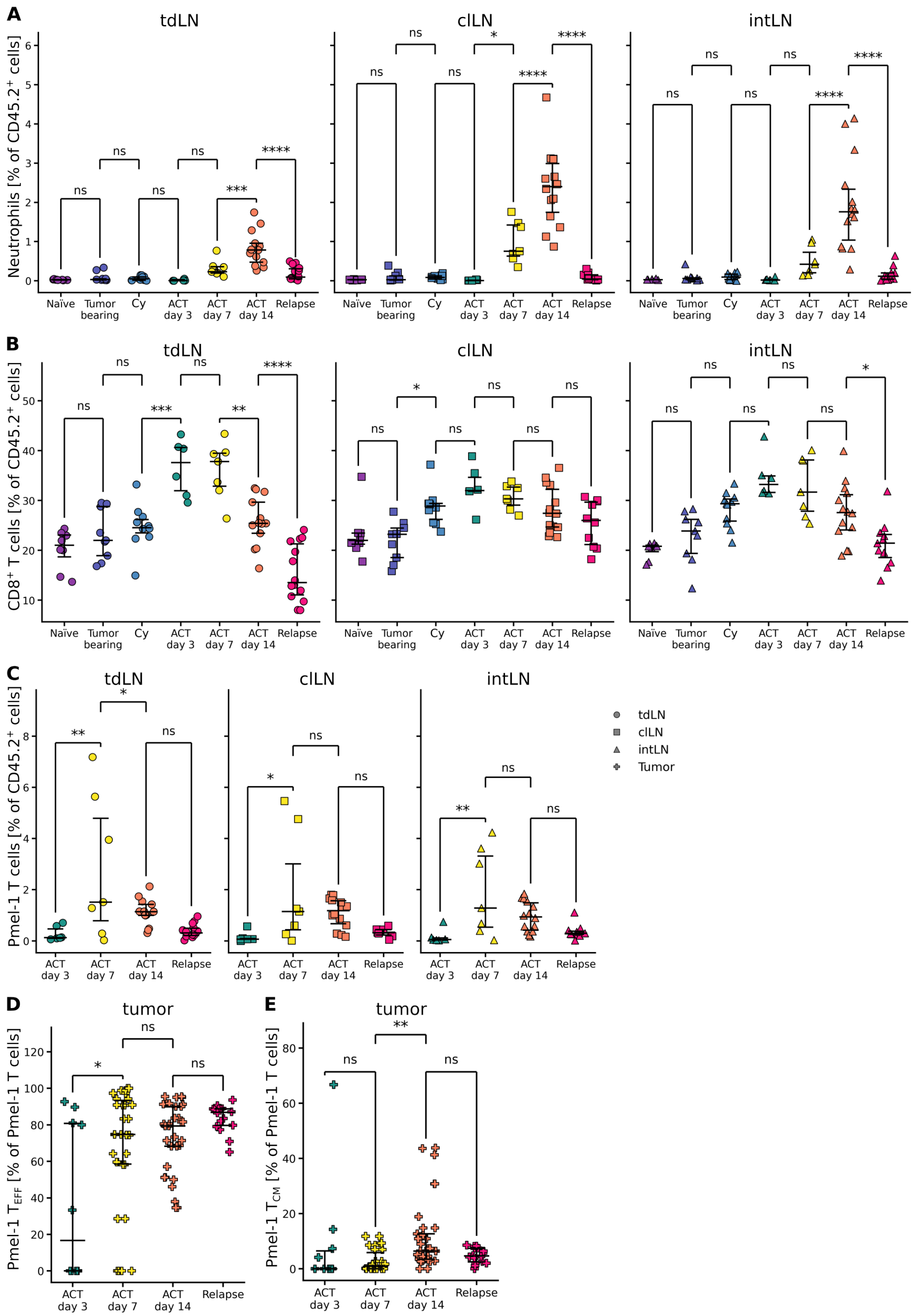


Figure S2: **Abundances of major cell types in the lymph nodes and tumor between conditions. (A-C)** Neutrophils, $CD8^+$ T cells and Pmel-1 T cells in the tumor draining (left), contra-lateral (middle) and intermediate (right) lymph nodes. **(D, E)** Relative abundances of effector $CD8^+$ T cell ($T_{EFF}$) and central memory $CD8^+$ T cell ($T_{CM}$) in the tumor. Unpaired tests **(A-E)** performed using ANOVA with Tukey's HSD for family-wise error correction. All data was obtained using flow cytometry.

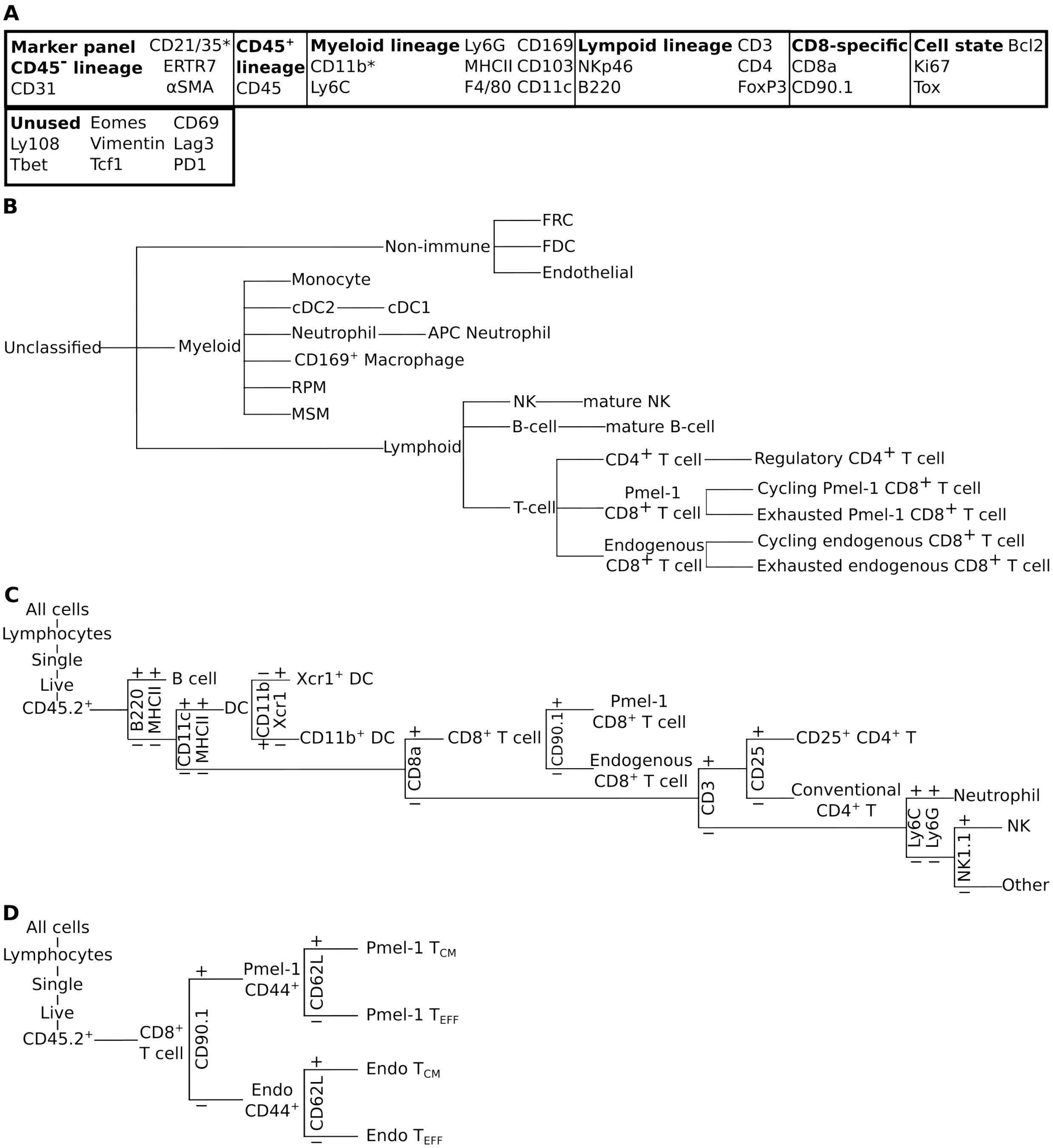


Figure S3: **Overview of phenotyping strategies.** **(A)** Multiplex immunofluorescence imaging marker overview. 22 markers where used for phenotyping. A * indicates a marker was used for phenotyping more than one cell type. **(B)** Multiplex immunofluorescence imaging phenotyping tree. Exhausted Pmel-1 $CD8^+$ T cells were not found. **(C)** Flow cytometry pan-immune panel. **(D)** Flow cytometry $CD8^+$-specific panel.

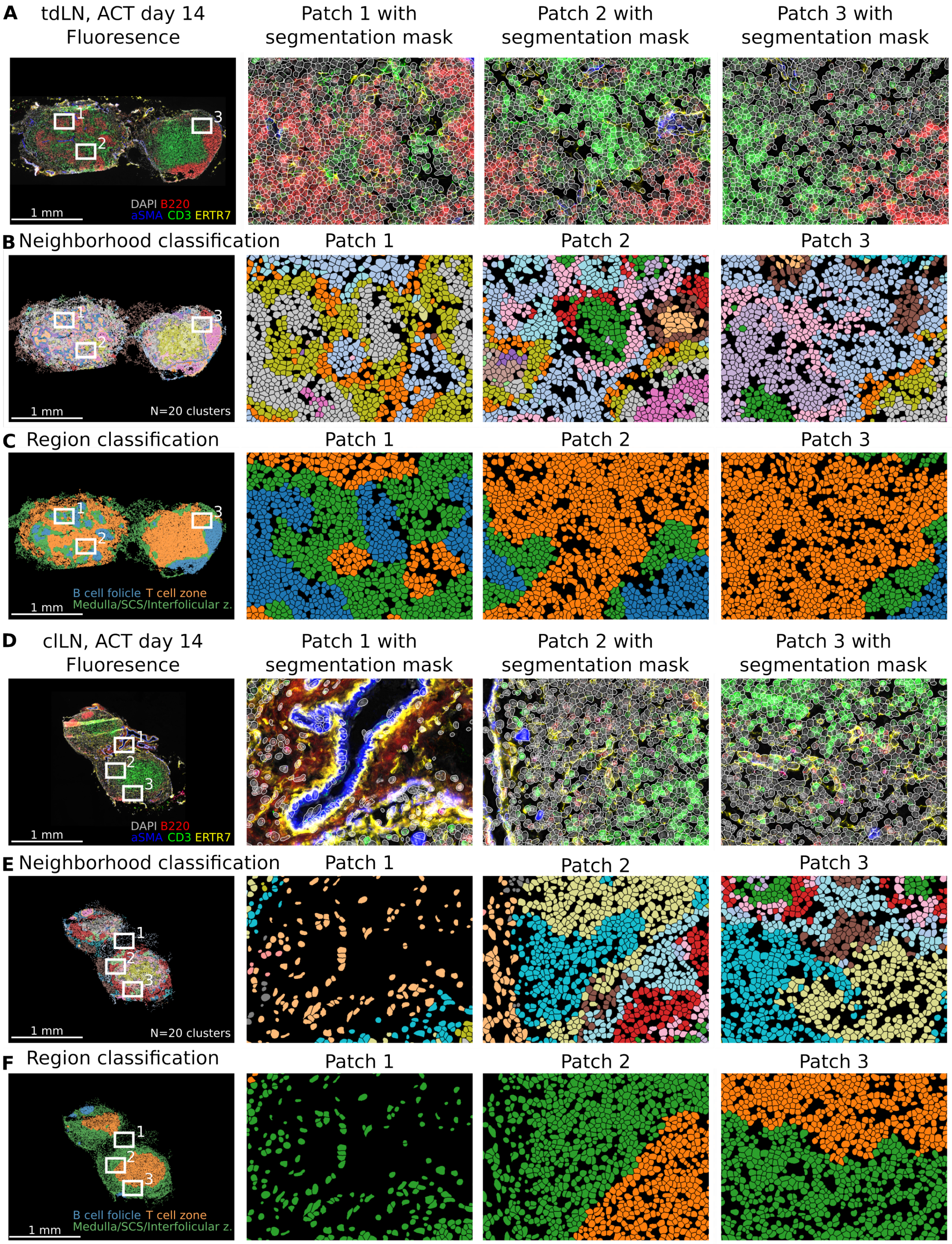


Figure S4: **Additional illustration of segmentation, niche and region classification in multiplex immunofluorescence imaging analysis (A, D)** tumor-draining lymph node (tdLN) **(A)** and contralateral lymph node (clLN) **(D)** at adoptive T cell therapy (ACT) day 14 with segmentation masks for three image patches. Selected markers: DAPI (grey), B220 (red), aSMA (blue), CD3 (green), ERTR7 (yellow). **(B, E)** Lymph node and zooms from **(A, D)** with neighborhood classification based on $k = 50$ nearest neighbors and $n = 20$ clusters. **(C, F)** Lymph node and zooms from **(A, D)** with smoothed region classification. B cell follicle (blue), T cell zone (orange) and Medulla/subcapsular sinus (SCS)/Interfollicular zone (green).

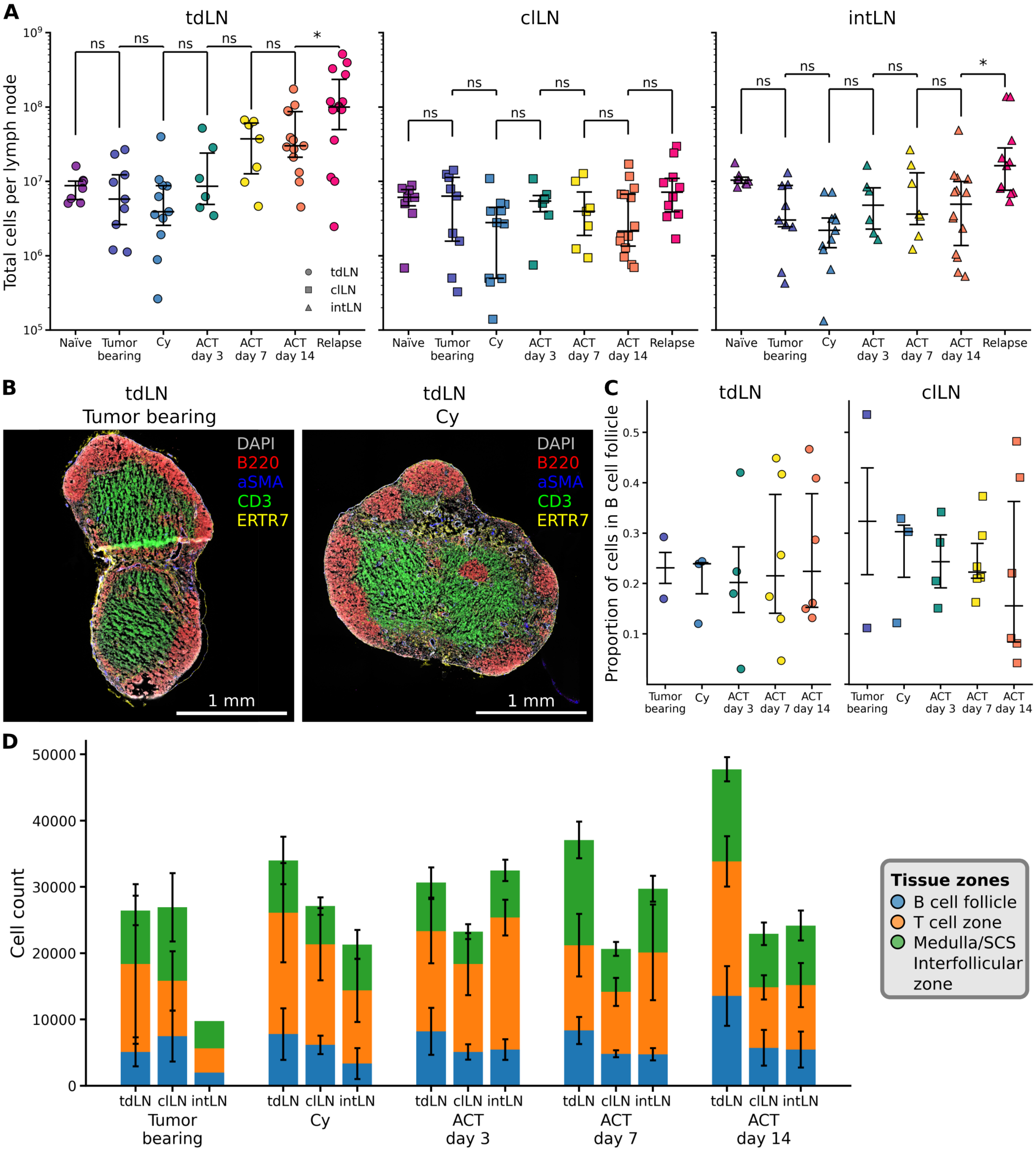


Figure S5: **Size differences, architectural similarities and compositional changes in the lymph nodes.** **(A)** Total live cell counts per lymph node in flow cytometry data, compared between conditions. **(B)** Representative tdLN lymph nodes in the tumor bearing and cyclo conditions. Selected markers: DAPI (grey), B220 (red), aSMA (blue), CD3 (green), ERTR7 (yellow). **(C)** Relative size of the B cell follicle compartment in the tdLN and clLN. **(D)** Averaged sizes of the lymph nodes in the imaging data, split by compartment. Error bars indicate standard error of the mean. Unpaired tests **(A, C)** performed using ANOVA with Tukey's HSD for family-wise error correction. Non-significant associations in **(C)** are not shown.

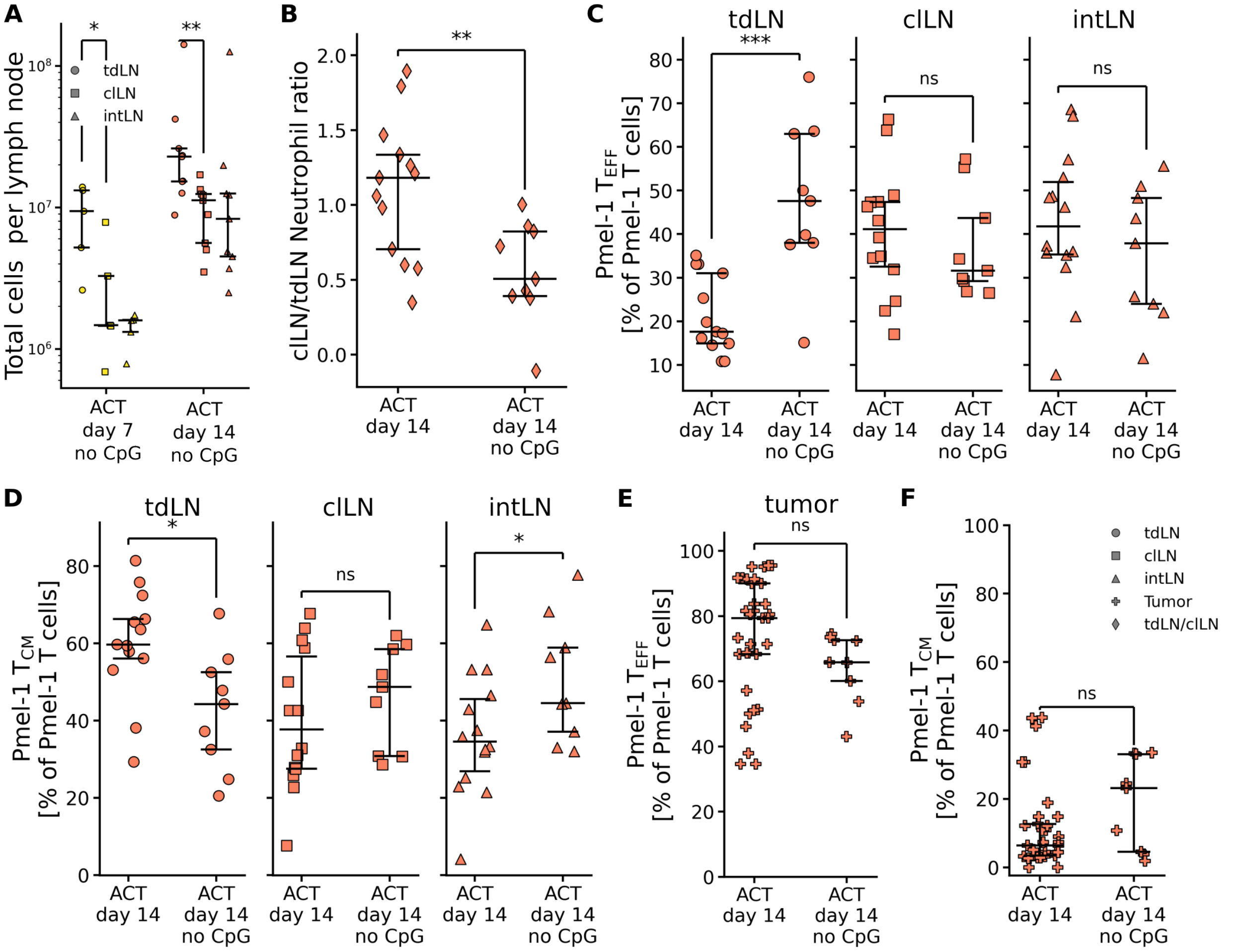


Figure S6: **Additional comparisons on the omission of innate immune stimulation in ACT day 7 and 14.** **(A)** Lymph node sizes with and without CpG/Poly(I:C) treatment on day 7 and 14. **(B)** The log $\frac{\text{clLN}}{\text{tdLN}}$ of neutrophils as a percentage of $CD45^{+}$ cells with and without CpG/Poly(I:C) treatment at day 14. **(C, D)** $T_{EFF}$ and $T_{CM}$ with and without CpG/Poly(I:C) treatment at day 14, per lymph node. **(E, F)** $T_{EFF}$ and $T_{CM}$ in the tumor. Paired tests **(A)** performed using a ratio-paired T test. Unpaired tests **(B-E)** performed using ANOVA with Tukey's HSD for family-wise error correction. All data was obtained using flow cytometry.

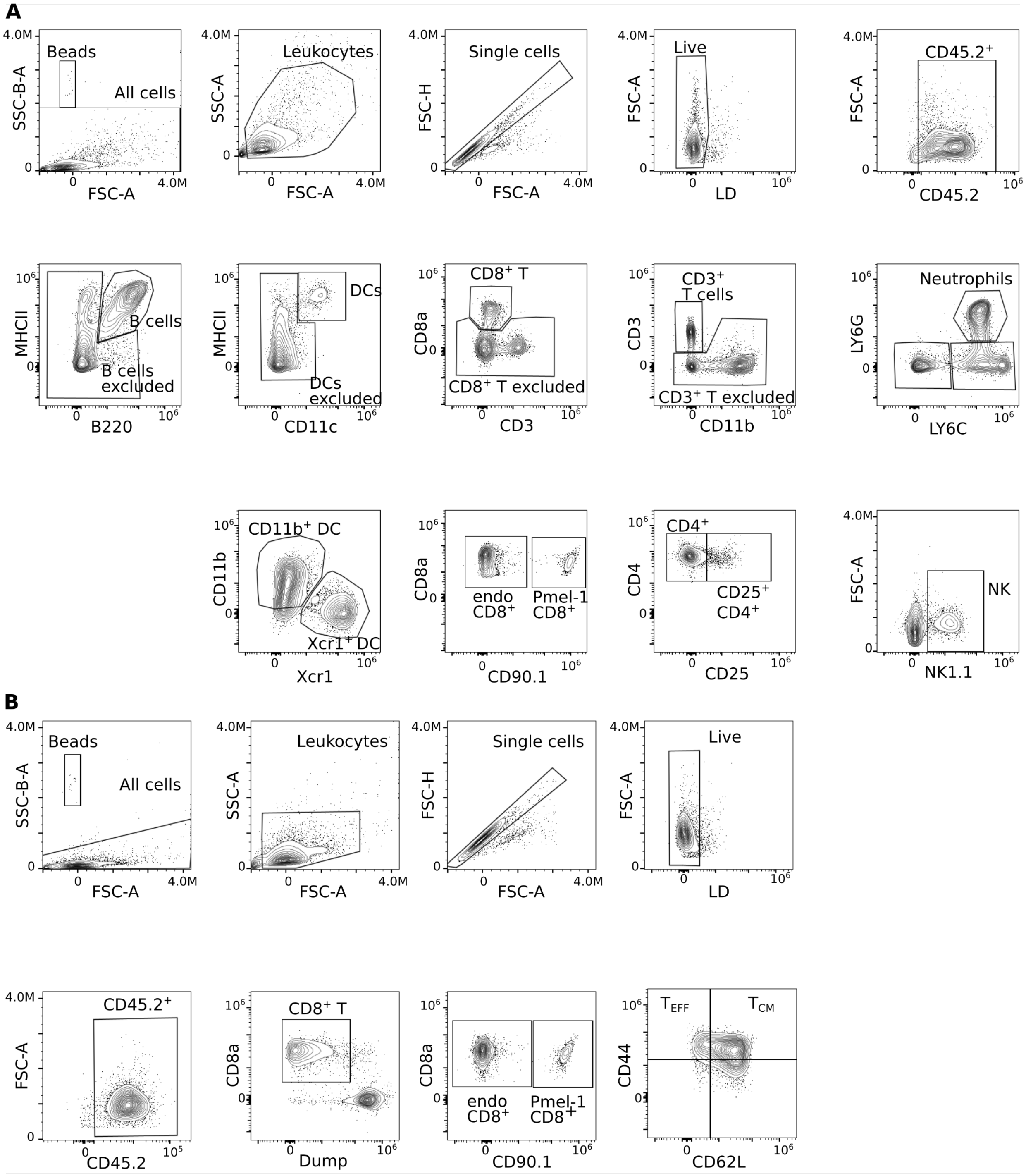


Figure S7: **Details of flow cytometry gating strategy.** **(A)** pan-immune panel **(B)** CD8$^+$-specific panel.
